# LDLR variant classification through activity-normalized prime editing screening

**DOI:** 10.64898/2025.12.16.694467

**Authors:** Phillip J. Zhou, Minja Velimirovic, Tian Yu, Vojislav Gligorovski, Nicolas Mathis, Jing Zhao, Quang Vinh Phan, Felicitas Vogd, Jayoung Ryu, Qisheng Pan, Atharva Tyagi, David B. Ascher, Gerald Schwank, Luca Pinello, Christopher A. Cassa, Richard I. Sherwood

**Author notes:** These authors contributed equally (Phillip J. Zhou, Minja Velimirovic). Corresponding authors name, email address, and address: Christopher Cassa, Division of Genetics, Brigham and Women’s Hospital and Harvard Medical School, 77 Avenue Louis Pasteur, Boston MA 02115, Richard Sherwood, Division of Genetics, Brigham and Women’s Hospital and Harvard Medical School, 77 Avenue Louis Pasteur, Boston MA 02115.

## Abstract

**Background:** Inherited variants in the LDL receptor (*LDLR*) gene are the most common cause of familial hypercholesterolemia (FH), significantly increasing coronary artery disease risk. Early identification of pathogenic LDLR variants enables prompt intervention with lipid-lowering therapies; however, the majority of *LDLR* variants observed in the population have uncertain or absent clinical classifications, limiting the potential to improve clinical management.

**Methods:** We developed an innovative, activity-normalized prime editing screening pipeline to measure the impact of 5,184 *LDLR* coding variants on LDL-cholesterol (LDL-C) uptake. Through pairing a genotypic outcome reporter with every prime editing guide RNA (pegRNA), we adjust phenotypic measurements to account for variable editing efficiency, extending activity normalization to prime editing for the first time at this scale. Further, we use a statistical estimation approach that leverages measurements for all missense variants at a given position to denoise the resulting scores.

**Results:** We show that prime editing-mediated reporter editing correlates with endogenous variant installation frequency, allowing activity normalization to improve imputation of *LDLR* variant effect. Our optimized prime editing assay identifies a broad, continuous spectrum of variant functional effects. We achieve robust separation of pathogenic *vs.* benign ClinVar variants and concordance between experimentally derived functional scores and LDL-C levels measured in UK Biobank participants. Further, when calibrating the strength of evidence provided by this functional screening data to align with the ACMG/AMP variant interpretation guidelines, and integrating additional sources of evidence, a majority of currently unclassified rare *LDLR* variants meet evidence thresholds for reclassification. We use the broad coverage of this screen to gain insight into how apolipoproteins bind to LDLR. In particular, we identify and characterize rare LDLR variants that enhance LDL-C uptake through increased interaction with apolipoprotein B. Finally, we compare prime editing-based functional scores with those derived from recent base editing and cDNA-based LDLR variant screens, showing that these approaches all show robust correlation with clinically observed LDL-C levels and computational scores, while prime editing identifies candidate splice-altering coding variants that are not modeled by cDNA screening.

**Conclusions:** Altogether, our approach demonstrates the power of prime editing to significantly improve understanding of how variants in *LDLR* impact function and contribute to FH.

## Introduction

The low-density lipoprotein receptor (LDLR) is the cell surface receptor responsible for removal of low-density lipoprotein cholesterol (LDL-C) from circulation^1^. Rare *LDLR* coding variants influence LDL-C levels, and deleterious variants are the primary cause of familial hypercholesterolemia (FH), a condition characterized by exceptionally high LDL-C levels and an increased risk of premature cardiovascular disease^2^. As coronary artery disease (CAD) risk depends on lifelong serum LDL-C exposure^3,4^, early diagnosis and treatment is key to risk reduction in FH patients^5,6^.

Nonetheless, a substantial number of *LDLR* variants lack definitive clinical interpretation. Of the 1,744 LDLR missense variants recorded in the ClinVar database^7^, over half are either designated as variants of uncertain significance (VUS) or have conflicting assessments of pathogenicity. Moreover, ClinVar-annotated variants represent only a subset of *LDLR* coding variants observed in the population, there are hundreds of additional rare *LDLR* missense variants observed in whole exome and whole genome sequencing in the UK Biobank (UKB)^8^ that have not yet been submitted to ClinVar. Thus, at present, a substantial number of patients with *LDLR* variants cannot receive potentially actionable information about the consequences of their genotype. Enhancing understanding of *LDLR* variant impacts could improve the early diagnosis and treatment of individuals with FH.

One reason that so many variants have uncertain classifications is that most *LDLR* variants are too rare to be confidently classified using human genetics evidence alone, and additional evidence has not been sufficient to reach a classification. Cellular LDL-C uptake assessment provides a scalable approach to functionally characterize the effects of *LDLR* variants. Prior efforts have used microscopic^9,10^ and flow cytometric^11,12^ measurement of fluorescent LDL-C uptake to improve classification of *LDLR* variant effects. However, these prior studies have limitations that motivate further efforts to profile *LDLR* variant effects. Thormaehlen *et al*^9^, Islam *et al*^10^, and Tabet *et al*^12^ evaluated LDL-C uptake capacity of cells engineered to overexpress cloned *LDLR* variant constructs. Overexpression of proteins can lead to artifactual findings^13^, impeding the ability to detect variants that alter protein stability and accentuating dominant negative effects^14^. Additionally, such screens cannot evaluate variants that impact splicing or gene regulatory activity. Ryu *et al*^11^ used CRISPR base editing^15,16^, in which a Cas9-nickase enzyme is fused to a deaminase enzyme to install transition variants into endogenous genes. While this approach introduced variants into endogenously expressed LDLR, base editing is inherently limited in the scope of variants that can be installed. This study was only able to collect functional data on 12% of LDLR variants observed in the UKB cohort.

CRISPR prime editing (PE), in which a Cas9-nickase is fused to a reverse transcriptase (RT) enzyme^17^, offers a more versatile and precise method for endogenous introduction of genetic variants. Through encoding desired sequence replacements in an RT template appended to the guide RNA, referred to as a prime editing guide RNA (pegRNA), PE enables installation of all single-nucleotide variants and codon swaps. Prime editing screening has been shown to be a powerful tool to decipher variant effects in high-throughput cellular screens^14,18–23^.

Since its original introduction, PE efficiency has been improved by a series of advances in the design of the enzyme and pegRNA as well as in the interaction between PE and cellular hosts. Prime editors have been improved through codon optimization and appending peptides that increase translation efficiency^24,25^. PegRNA degradation has found to limit PE efficiency and has been enhanced by adding pseudo-knot structures to the end of pegRNAs as well as appending the prime editor enzyme with the La protein that stabilizes 3’ RNA ends^26,27^. PE was shown to be antagonized by cellular mismatch repair (MMR)^24,28^; performing PE in cellular models with deficient MMR improves efficiency. Finally, efforts combining high-throughput PE efficiency measurement and machine learning have led to tools such as PRIDICT/PRIDICT2.0 that predict the most efficient pegRNA for a desired edit^29,30^.

While these efforts have dramatically improved editing efficiency and reliability, PE remains limited by incomplete and variable installation efficiency. This variability complicates implementation of PE for massively parallel screening applications. CRISPR screens typically rely on using guide RNA abundance in phenotypically defined cell populations as a surrogate for genome edits, so the simplest implementation of PE screening would be to identify pegRNAs with enriched or depleted representation in phenotypically defined populations. In the presence of variable editing, this approach is inaccurate, as inefficient pegRNAs are erroneously interpreted as phenotypically inert. Our prior base editing screening work introduced activity normalized screening, in which each gRNA is paired with a reporter construct that contains the genomic target sequence of that gRNA^11^. Importantly, we developed a novel computational pipeline, BEAN, that uses a Bayesian generative model to normalize and deconvolve the phenotypic scores of target variants by using the relative frequencies and identities of genotypic outcomes collected from the target site reporter. Such an approach has yet to be applied to PE screening. Prior work has used a paired reporter to filter out pegRNAs with below-threshold editing efficiency^14^, but this work did not account for variable efficiency above this threshold.

Here, we introduce an activity-normalized prime editing (ANPE) platform and use it to evaluate *LDLR* variants at an unprecedented scale through deep mutational scanning. We benchmark the activity-normalized screening approach by correlating reporter and endogenous editing activity across 1,646 variants. We use ANPE to measure the impact of a total of 5,184 *LDLR* coding variants on LDL-cholesterol (LDL-C) uptake across two screens. The functional scores derived from these screens robustly distinguish positive control (stop-gain, ClinVar pathogenic) *vs.* negative control (synonymous, ClinVar benign) variants, associate with a range of computational variant pathogenicity metrics, and show concordance with LDL-C levels measured in UK Biobank participants. Using this wealth of functional data alongside computational and population evidence, we find that 74.2% of *LDLR* variants observed in the UK Biobank that currently lack a definitive classification meet evidence thresholds for reclassification. Further, we identify and characterize rare LDLR variants that enhance LDL-C uptake through increased interaction with apolipoproteins and show that such variants can rescue activity when co-installed with pathogenic missense variants. Finally, we compare scores from base editing-, prime editing-, and cDNA-based LDLR variant scans, finding that they provide broadly similar results, although prime editing flags candidate splice-altering variants as deleterious in contrast to cDNA scores and computational pathogenicity predictions. Altogether, this study introduces a novel genome editing platform to substantially improve *LDLR* variant classification and FH diagnostics.

## Methods

### LDLR pegRNA Library Design

We developed a computational pipeline to design pegRNAs for comprehensive mutational screening. The pipeline generated all possible pegRNAs for each desired amino acid swap, and this exhaustive set was subsequently used as input for PRIDICT/PRIDICT2.0^29,30^ to prioritize pegRNAs predicted to yield high-efficiency genome editing.

As a first step, a set of target amino acid swaps was generated. For the LDLR137-219 library, this set included all 21 amino acid swaps (missense, synonymous, and stop-gain) at positions 137-219 of LDLR. For the LDLR-FL library, this set included (i) four missense variants at all LDLR positions, optimized for imputation quality; (ii) all LDLR amino acid swaps in the ClinVar database and/or observed in UK Biobank whole exome sequencing cohort; (iii) 600 synonymous and 600 stop-gain variants across LDLR.

To optimize our missense variants for imputation quality, we designed the LDLR-FL screening library with the GLIDE pipeline (manuscript in preparation) to prioritize, at each codon, the four prime-editable coding substitutions expected to yield the lowest imputation error, or in other words, the most informative variants for predicting the effects of untested mutations. By supplementing these GLIDE-selected edits with all LDLR coding variants observed in ClinVar and the UKB, we ensure that the library provides direct functional readouts for known human alleles while supplying a highly informative scaffold for FUSE-based imputation of the remaining untested substitutions. This GLIDE-driven optimization (manuscript in preparation) couples empirical coverage of real-world variants with principled selection of additional edits to improve imputation accuracy using FUSE^31^.

The design process then generated an exhaustive list of candidate nucleotide changes and paired spacers [(variant, PAM) pairs] to fulfil each target amino acid swap. To do so, every possible codon swap that produces a target amino acid swap (up to 63 potential codon swaps per AA) was considered. Then, a set of candidate nucleotide changes for that codon swap was generated by considering all local PAMs. The pipeline searched for nearby PAM sites (“NGG,” in practice “GG”), located within 17 bp upstream (sense strand) or 17 bp downstream (antisense strand) of the target codon swap, a window that was extended to 35 bp in rare cases where proximal PAMs were not identified.

When a PAM was identified, the following design rules were applied to generate a (variant, PAM) pair that precluded retargeting by Cas9 after installation:

1. **Variant disrupts PAM/seed** – If the target codon swap alters at least one “G” in the PAM sequence or the seed sequence (the three nucleotides between the PAM and the nick site), thus preventing repeated editing of the same site, the codon swap is sufficient and the (variant, PAM) pair could consist of the variant only.
2. **Adjacent synonymous PAM/seed disrupting variant** – If the target codon swap does not disrupt the PAM or seed, introduce a synonymous variant to disrupt the PAM (top priority) or seed (second priority) such that the final sequence change include both the target codon swap and this nearby synonymous variant.
3. **Ensure Hamming distance >1** - To facilitate accurate sequencing-based readouts, we further require that the wild-type and variant sequences differ at a minimum of two nucleotides (Hamming distance > 1). If the above two steps do not naturally satisfy this condition, add an additional synonymous variant within 1–3 codons upstream of the target codon swap.

All nucleotide changes were constrained to lie within the coding sequence to avoid disrupting intronic splice sites. For target codon swaps for which multiple synonymous options exist that satisfy these rules equally well, all valid (variant, PAM) pairs were retained to maximize the likelihood of identifying a pegRNA with high predicted efficiency in PRIDICT/PRIDICT2.0. This approach produced, on average, ∼15 (variant, PAM) pairs per target codon swap. Because many amino acids can be encoded by multiple codons, there was an even larger number of (variant, PAM) pair candidates per target amino acid swap.

All (variant, PAM) pairs were formatted and run through PRIDICT (LDLR137-219)/PRIDICT2.0 (LDLR-FL). While PRIDICT/PRIDICT2.0 does not allow specification of the spacer to be used for a given submission, we filtered the output to retain the top-scoring pegRNA for which the spacer utilized the appropriate PAM in the input (variant, PAM) pair. Of the dozens of top-scoring pegRNAs for (variant, PAM) pairs corresponding to a given amino acid swap, we then selected the single pegRNA per target amino acid swap with the highest predicted HEK293T installation efficiency for inclusion in the final library.

This pipeline produced pegRNA candidates for the vast majority of desired amino acid swaps. Codons that spanned splice junctions were not included, as it was not possible to encode all desired codon swaps at these positions with short sequence replacements.

These top pegRNA candidates were then formatted for oligonucleotide library synthesis and cloning. The library structure was as follows:

TGTGGAAAGGACGAAACACCG[19-nt spacer] GTTTCAGAGCTATGCTGGAAACAGCATAGCAAGTTGAAATAAGGCTAGTCCGTTATCAACTTGAAAAAGTGGCACCGAGTCGGTGC [PBS/RT] CGCGGTTCTATCTAGTTACGCGTTAAACCAACTAGAATTTTTT [Reporter] [7-nt barcode] AGATCGGAAGAGCACACGTCT

Further filtering was performed for pegRNA inclusion in the LDLR-FL library. The encoded pegRNA library member with full target sequence was required to be <=300-nt, which excluded pegRNAs far away from PAMs which require long target reporters.

### Generation of HCT116 cell line for prime editing screening

An HCT116 cell line was constructed for high-efficiency lentiviral prime editing screening. First, a clonal HCT116 TASOR knockout line was generated as follows. Nucleofection mix was prepared by mixing 300 pmol TASOR sgRNA (synthesized by IDT as Alt-R CRISPR-Cas9 sgRNA; Spacer sequence 5’-CTCGTGGCTGAGGCTCTGGG-3’) with 100 pmol of SpCas9 (IDT Alt-R CRISPR-Cas9), 1.4 ul of 30% (v/v) glycerol, 16.4 ul Nucleofection solution and 3.6 ul supplement solution (SE Cell Line 4D-Nucleofector X Kit), and incubated for 15 minutes at room temperature. 400,000 trypsinized HCT116 cells were spun down at 300xg for 5 minutes, resuspended in nucleofection mix and pulsed in the Lonza Amaxa 4D-Nucleofector system with the EN-113 nucleofection program. Cells were recovered in McCoy’s 5A medium and kept in culture for 2 weeks until subcloning via limited dilution. Clones were verified for frameshift mutation in TASOR via PCR/Sanger sequencing and one clone was selected for further experiments.

HCT116 TASOR-KO cells were stably transfected with Tol2 transposase along with a Tol2 transposon-based prime editor vector (p2T_CAG_SpCas9PE2_P2A_mCherry_BlastR). Transfected HCT116 TASOR-KO cells were selected via blasticidin (10 ug/ml) and subsequently flow cytometrically sorted to enrich for cells with high mCherry signal.

### Prime editing screening workflow

The pegRNA libraries were cloned into pLenti U6_r2ad_ IN-PE2_PuroR (LDLR137-219)/ pLenti U6_r2ad_ IN-PE2-SSB_PuroR (LDLR-FL). Libraries were amplified using NEBNext Ultra II Q5 Master Mix, cloned using NEBuilder HiFi DNA Assembly Master Mix and electroporated into Endura electrocompetent cells (Biosearch Technologies) for propagation. Lentivirus was produced in HEK293T cells (ATCC CRL-3216) for each library using TransIT-Lenti transfection reagent, titered, and incubated with 3.91x 10^6 HCT116 TASOR-KO PE2-mCherry cells per replicate at a multiplicity of infection of 0.3-0.5. After 24 h, medium containing lentivirus was removed and fresh McCoy medium. After another 48 h, medium with 500 ng/mL puromycin was added and cells underwent selection for the next 5-7 days, splitting as needed.

After complete selection defined by complete death of a concurrent untransduced HCT116 control, cells were started on a 2 day LDL uptake protocol. On day 0, library transduced cells were plated onto 15 cm plates, 39.1 million cells per plate. On the evening of day 1, growth media was replaced with Opti-MEM (Thermo Fischer Scientific) to induce overnight serum starvation. On the morning of day 2, Opti-MEM with 2.5 ug/mL BODIPY FL-LDL (Thermo Fisher Scientific) were incubated with the cells for 4 hours. After this incubation period, cells in plates were trypsinized, stained with 50 ng/mL DAPI and sorted based on BODIPY-LDL fluorescence levels into four bins (top 20%, top 20-40%, bottom 20% and bottom 20-40%). Genomic DNA (gDNA) was isolated from each sorted population as well as from an unsorted bulk population via PureLink Genomic DNA Mini Kit (Thermo Fischer Scientific) and prepared for Next Generation Sequencing.

### Nextgen sequencing library preparation and sequencing

gDNA collected from each population was used as the input for PCR1 aimed at amplifiying the integrated construct spanning the pegRNA and reporter sequence as well as adding different inline PE1 barcodes to specific samples for downstream analysis. Total gDNA input per sample was 20 ug and NEBNext Ultra II Q5 Master Mix was used for all PCR reactions. Following PCR1, reactions were individually PCR purified using a standard QIAquick PCR Purification Kit. Next, a qPCR2 was performed from 0.25 μl of each purified sample in a 15-μl qPCR reaction to determine the optimum number of cycles for PCR2, and the products of qPCR2 were run on a gel to confirm lack of primer dimers. PCR2 cycle counts were chosen to be two to three cycles less than the qPCR2 Ct for the corresponding sample, with the minimum number of PCR2 cycles being five. To set up PCR2, half of each sample’s purified PCR1 product was then used with NEBNext Ultra II Q5 Master Mix. After PCR2, samples were PCR purified again and were then run on a 2200 Agilent TapeStation to quantify the product. Samples were then pooled based on their molarity, and the pool was purified to remove unwanted products using SPRIselect beads (Beckman Coulter). After bead purification, the pooled sample was sequenced using the Element AVITI sequencer with paired-end sequencing and 150 nucleotides for read 1 and 150 nucleotides for read 2.

### Estimating variant effects using the BEAN-FUSE pipeline

We made use of the BEAN-FUSE pipeline to improve the estimation of variant functional effects and to impute effects of variants that had not been screened^11^. BEAN provides a create-screen mode that allows users to construct a ReporterScreen directly from flat count tables, bypassing the count-samples module, which is designed for processing base-editing data that generates multiple edited alleles. In prime editing, each pegRNA introduces a single, deterministic edit, enabling BEAN to analyze prime-editing data by treating each pegRNA as an independent variant. We used the create-screen mode to build a ReporterScreen directly from our pegRNA library and count data, followed by the standard BEAN workflow to estimate variant effect sizes.

FUSE makes use of related measurements within and across experimental assays, namely an amino acid substitution matrix derived from 115 deep mutational scanning datasets, to jointly estimate variant impacts^31^.After functional scores had been estimated by the BEAN pipeline, the full set of scores was processed by FUSE, which first collectively estimates the mean functional effect per amino acid residue position within the assay using shrinkage estimation. FUSE then makes estimates for individual allelic variants within the amino acid residue position, based on a functional substitution matrix derived from deep mutational scanning data across many genes. The result is a full set of estimated variant functional effects for both (1) the original variants screened in the assay and (2) other possible variants that were not screened but fell within amino acid residues that had variants covered in the screen.

### UK Biobank variant and phenotype processing

The UK Biobank^32^ is a large prospective cohort comprising over 500,000 individuals aged 40–69 years when recruited, between 2006 and 2010. From 469,803 participants with available whole-exome sequencing data, we included 443,353 individuals with valid low-density lipoprotein cholesterol (LDL-C) measurements. We identified rare (allele frequency <1:1,000) variants in the canonical LDLR coding sequence. To minimize confounding from additional genetic contributors to lipid levels, participants carrying homozygous variants, multiple rare variants within *LDLR*, or any rare variant in *APOB*, *PCSK9*, or *ANGPTL3* were excluded. After applying these filters, 8,892 participants harboring 717 unique *LDLR* coding variants were retained for analysis.

Exon coordinates for *LDLR*, *APOB*, *PCSK9*, and *ANGPTL3* were defined using MANE^33^ transcript annotations. Exome sequencing of UK Biobank participants was performed as previously described^34^ and analyses were conducted on the UK Biobank Research Analysis Platform (https://ukbiobank.dnanexus.com). Gene-level variant call files (VCFs) were extracted from the joint-called whole-exome sequencing pVCFs using BCFtools (v1.15.1) and Swiss Army Knife (v4.9.1), followed by normalization to flatten multi-allelic sites and alignment to the GRCh38 reference genome. Variants located in low-complexity regions, segmental duplications, or other genomic regions known to be difficult for next-generation sequencing alignment or variant calling (as defined by the National Institute of Standards and Technology Genome in a Bottle Consortium^35^) were excluded. We further filtered variants with an alternate allele frequency greater than 0.1% in the UK Biobank cohort, variants missing genotype calls in more than 10% of samples, and those absent from the cohort. To minimize coverage-related biases between sequencing phases, only variants with at least 90% of called genotypes supported by a minimum read depth of 10 were retained. Noncoding variants (that were not canonical splice sites) were excluded from all downstream analyses.

Phenotypic data related to coronary artery disease (CAD) and myocardial infarction (MI) were aggregated from hospital records (primary or secondary diagnosis), death registries (primary or secondary cause of death) and self-reported data. Patient-level LDL-C values were ascertained from UKB data files. Estimated untreated LDL-C levels obtained using adjustments for lipid-lowering therapies were used in analyses, as described in *Ryu et al.*^11^.

### ClinVar variant classifications

ClinVar variant classifications were obtained from the tab-delimited ClinVar^7^ release on 25 July 2025. *LDLR* coding variants were cross-referenced against this database, where variants annotated as *pathogenic*, *likely pathogenic*, or *pathogenic/likely pathogenic* were collectively labeled as P/LP, and those annotated as *benign*, *likely benign*, or *benign/likely benign* were collectively labeled as B/LB. All remaining variants retained their ClinVar annotations of variants of uncertain significance (VUS) or conflicting classifications of pathogenicity, and variants not presented in ClinVar were labeled as unclassified.

### Statistical Comparison of FUSE Score Distributions

Statistical comparisons were performed using two-sided non-parametric Mann–Whitney U tests. Only observed variants presented in the LDLR-FL functional screen were included in the analysis. FUSE score distributions were compared across ClinVar variant categories (“B/LB”, “VUS/conflicting”, and “P/LP”), between UK Biobank participant groups defined by LDL-C levels (carriers with LDL-C ≥ 190 mg/dL versus those with LDL-C < 190 mg/dL), and between participants with CAD or without CAD in the UK Biobank. Statistical significance is indicated as * (p < 0.05), ** (p < 0.01), *** (p < 0.001), and **** (p < 0.0001).

### Calibrating the strength of functional screening evidence in alignment with clinical guidelines

The American College of Medical Genetics and Genomics and the Association of Molecular Pathology have developed Sequence Variant Interpretation (ACMG/AMP SVI) guidelines that describe standardized evidence types and strengths that are useful for variant classification. We calibrated our functional and computational evidence using the statistical framework described by Pejaver *et al.*^36^. ClinVar *P/LP* and *B/LB* variants were used as labeled reference data, with a prior probability of 0.201, corresponding to the probability that a variant is classified as P/LP in ClinVar given that it is observed in a UKB participant with a rare *LDLR* coding variant. The model estimates the maximum likelihood posterior probabilities that a variant is benign or pathogenic, and these posterior probabilities were subsequently used to assign quantitative evidence strength scores to individual variants.

### Developing ACMG/AMP SVI evidence for classification for variants within LDLR

To identify how many variants which are uncertain (VUS) or unclassified within ClinVar which have sufficient evidence to be classified as P/LP or B/LB, we considered multiple evidence sources and converted them to the semi-quantitative point system adaptation of the ACMG/AMP sequence variant interpretation (SVI) framework^37^. We integrated four forms of evidence which we were able to electronically assemble without a manual curated review of clinical variant summaries, including computational scores (AlphaMissense)^38^, population enrichment of disease at the variant level^39^, clinical context from previously classified variants within the same amino acid residue, and the functional evidence developed using activity normalized prime editing.

Consistent with the Bayesian adaptation of the ACMG/AMP SVI framework^37^, for each variant, each form of available evidence was assembled, and the appropriate number of points were added or subtracted, given the strength of each form of evidence. AlphaMissense computational scores were applied at strength levels developed using the calibration thresholds for evidence types PP3 or BP4 developed above. Similarly, prime editing functional scores were also applied at the appropriate strength levels for PS3 or BS3, based on the thresholds calculated above. Population evidence, defined as the statistical enrichment of cases over controls in a population cohort (PS4) was applied consistent with thresholds developed using UKB participants with variants in *LDLR*^39^. Contextual evidence from previously classified pathogenic variants was applied in the following manner, developed by the Tocayo pipeline^39^: PVS1 (+8 points) for LOF variants, PS1 (+4 points) when a variant has a colocated P/LP variant encoding the same substitution and -4 points when a variant has a colocated B/LB variant encoding the same substitution (note that the benign equivalent is not an established ACMG/AMP criterion, but has been analyzed), and PM5 (+2 points) when a variant has a colocated P/LP variant encoding a different substitution and -2 points when a variant has a colocated B/LB variant encoding a different substitution (note that the benign equivalent is not an established ACMG/AMP criterion, but also has been analyzed.)

The final ACMG evidence score for each variant was computed as the sum of four evidence components: functional, computational, population, and clinical diagnostic evidence. Variants were grouped by total ACMG score to define categories corresponding to Benign (≤−7), Likely Benign (≤−1), Variant of Uncertain Significance (0-5), Likely Pathogenic (≥6), and Pathogenic (≥10) classes.

### Validation of individual variants

To generate lentiviral plasmids for doxycycline-inducible overexpression of LDLR in HepG2 or K562 cells under the control of a TRE (tetracycline-responsive element) promoter, we subcloned the lentiTet LDLR-mCherry fusion BlastR plasmid to install individual gain-of-function (GOF) or loss-of-function (LOF) variants, and plasmids containing LDLR-mCherry fusion BlastR with individual variants were themselves subcloned to generate plasmids with concurrent GOF and LOF variants. Plasmids were transformed into NEB Stable Competent E. coli for propagation, lentivirus was produced in HEK293T cells (ATCC CRL-3216) for each construct of interest, where wells were seeded in 6-well format with 3.84 x 10^5 cells per construct. After 48 hours, DMEM medium containing lentivirus was removed and fresh DMEM/RPMI medium with 500 ng/mL puromycin was added. Cells underwent selection for the next 5-7 days and were split as needed.

After complete selection as defined by complete death of a concurrent untransduced HepG2 or K562 control, cells were started on an LDL uptake protocol. For HepG2 cells, on day 0, cells were split, counted, and replated in 96-well format at 5×10^4 cells per well. For the GOF/LOF experiment, 3.75 x 10^4 cells were plated in each well along with 1.25 x 10^4 untransduced HepG2 cells. On the afternoon of day 1, DMEM medium was removed and replaced with Opti-MEM + 2 ug/mL doxycycline to induce the expression of LDLR-mCherry. On the morning of day 2, 2.5 ug/mL BODIPY™ FL-LDL (Thermo Fisher Scientific) and Opti-MEM were added and incubated with the cells for 4 hours. After 4 hours have elapsed, cells in plates were trypsinized and analyzed via flow cytometry.

For K562 cells, on the afternoon of day 0, 6 x 10^5 infected cells and 2 x 10^5 uninfected K562 cells were plated per each construct in Opti-MEM + 2 ug/mL doxycycline. On the morning of day 1, cells were centrifuged and plated at 1 x 10^5 cells per well in 96-well format. 1 hour and 30 minutes before preparing for flow cytometry, Pitstop 2 or DMSO was added to cells such that the overall concentration was 20 uM per well, and 1 hour before, BODIPY FL-LDL was added to a concentration of 2.5 ug/mL per well. Cells were then centrifuged and submerged in FACS SM containing DMSO if they were previously treated with DMSO and Pitstop 2 if treated with Pitstop 2, and then analyzed via flow cytometry. A gating strategy was applied to capture cells with high mCherry and low mCherry, and the log2 fold change was calculated between the mean signal in the FITC channel for the mCherry^high^ and mCherry^low^ population. The unpaired Welch t-test was applied to compare log2(mCherry^high^FITC/ mCherry^low^FITC) between samples.

### Structural analysis

Protein structures were visualized using PyMOL (version 3.1.4.1). Wild-type atomic interactions between residues were calculated with Arpeggio. Mutant structures were generated using MODELLER 10.7^40,41^, and their atomic interactions were calculated using the same approach. To generate predictions for the LDLR-APOB interaction, we used the structure of APOB bound to the class A repeats of LDLR (PDB: 9bde) as reported by Reimund *et al*. All structures besides the relevant chains (chain A, the APOB β-belt, and chain R, the class A repeats of LDLR), such as legobodies, were removed from the structure prior to analysis.

Protein stability was calculated using FoldX (version 4) and DDMut. To assess LDLR stability upon point mutations, the extracellular domain of LDLR (PDB: 1n7d) as reported by Rudenko *et al*. was first repaired using the *RepairPDB* command, followed by saturation mutagenesis with *BuildModel*.

In addition, DDMut-PPI was used to calculate change of binding ΔΔG values upon point mutations using the same APOB–LDLR structure (PDB: 9bde) as input.

Molecular interactions from Arpeggio were visually represented using the same color palette and style as used in Ryu et al 2024: hydrophobic interactions are depicted in ‘forest’, polar interactions are depicted in ‘orange’, carbonyl interactions are depicted in ‘blue’, hydrogen bonds are depicted in ‘red’, aromatic ring interactions are depicted in ‘pale green’, undefined interactions are depicted in ‘cyan’, metal-ligand coordination bonds are depicted in ‘purple’, and ionic interactions are depicted in ‘yellow’. Dashed lines depict clashes and van der Waals clashes, and all other interactions are represented with solid lines. Proximal interactions predicted from Arpeggio were not displayed.

### AlphaGenome analysis of splicing variants

To measure the change in splicing probability associated with each variant, we utilized the AlphaGenome API. We fetched the location of the LDLR transcript using a GENCODE-annotated GTF file, and resized it to a width of 1,048,576 base pairs centered around the LDLR transcript, which we call the sequence_interval. We defined ism_intervals, which are sections of DNA that we mutated contained within the larger sequence_intervals, as the 18 exons of LDLR, padded by 100 nucleotides on each side.

We used the CenterMaskScorer from the AlphaGenome variant scorer list, set the requested_output type to SPLICE_SITE_USAGE, which calculates the probability a given nucleotide is used as a splice site, and the aggregation_type to DIFF_MEAN, which calculates the difference of the mean values across all residues in the mask between the alternate and reference sequences. This pipeline ran for all 18 exons of LDLR.

## Results

### Developing activity-normalized prime editing screening

To establish activity normalized prime editing (ANPE), we asked whether a target reporter construct adjacent to the pegRNA in a lentiviral vector provides a faithful surrogate of endogenous editing efficiency. To test the concordance of reporter *vs.* endogenous prime editing at scale, we chose to perform saturation mutagenesis in a contiguous region of 83 amino acids in *LDLR* exon 4. The length of this region allows for next-generation sequencing (NGS) of this endogenous genomic region from a single amplicon. We selected LDLR residues 137-219, which span a highly conserved and functionally critical stretch encompassing LDLR class A repeats 3-5^42^.

We developed a computational framework to generate pegRNAs and paired reporters for activity-normalized prime editing variant scanning (**Figure S1, Methods**). Given a list of desired amino acid swaps, this pipeline designs an exhaustive set of candidate nucleotide changes for each variant to be installed that meet several design rules. First, the nucleotide change must ablate the PAM or seed sequence of a SpCas9 spacer, which has been shown to enhance DNA editing while minimizing indels^29^. If the variant itself does not ablate the PAM or seed, an adjacent synonymous variant is added to accomplish this goal. Second, the candidate genotype must contain two or more nucleotide swaps compared to the wild-type sequence (Hamming distance > 1) for clear distinction of edited outcomes *vs.* PCR and NGS errors. These candidate genotypes were used as input to PRIDICT^29^, and one pegRNA predicted to yield highest efficiency editing for each desired variant was selected for inclusion in a library paired with a reporter that included the spacer through to the end of the RT template.

To construct a saturation mutagenesis library for LDLR residues 137-219 (LDLR137-219 library), we sought to install all 21 possible amino acid swaps (19 missense, one synonymous, and one stop-gain variant) at each of these 83 residues. Our pipeline generated PRIDICT-optimized pegRNAs and reporters for 1738 of the 1743 variants (it was not possible to design pegRNAs with sufficient Hamming distance for the remaining variants) from an average of 26 candidate genotypes per desired variant (**Figure S1B**). We ordered a 300-nt oligo library with these pegRNAs and reporters embedded in an e-pegRNA template with a barcode following the reporter to facilitate disambiguation (**Figure 1A, Supplemental Table 1**). For 94 variants, the 300-nt oligo length was insufficient to include the full reporter, such that our library enabled reporter evaluation for 1646 pegRNAs.

**Figure 1.**
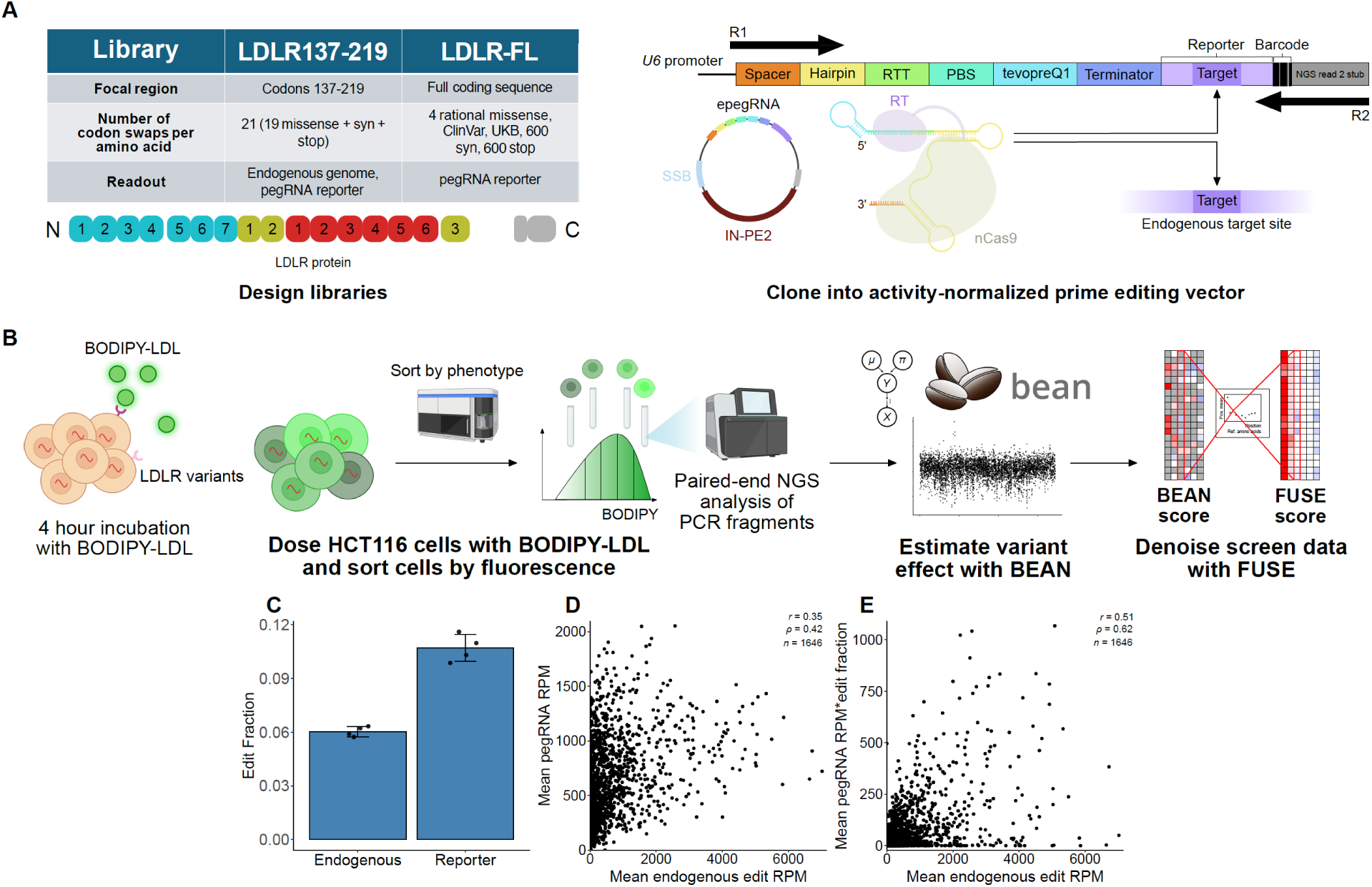
Activity-normalized prime editing screening (ANPE) pipeline and benchmarking. **A**, Schematic of design of LDLR137-219, LDLR-FL libraries, and the reporter system used for activity normalization. **B**, Schematic of the ANPE pipeline and downstream computational analysis. **C**, Overall prime edited allele fractions in unsorted LDLR137-219 endogenous and reporter replicates. **D-E**, Scatterplots of mean reporter pegRNA reads per million (RPM, **D**) or the product of mean reporter pegRNA RPM and mean reporter edit fraction (**E**) *vs.* mean endogenous edit RPM across unsorted LDLR137-219 replicates. *r*, *ρ*, Pearson, Spearman correlation coefficient.

To benchmark activity-normalized prime editing, we performed screens in HCT116 cells, which have been found to show high prime editing efficiency due to their lack of functional mismatch repair caused by bi-allelic MLH1 mutation^23,29^. As LDLR activity assays have been benchmarked in a range of cell types^9,12,43,44^ and HCT116 express LDLR robustly, we reasoned that HCT116 would provide a suitable cellular model for LDLR function. We engineered HCT116 to stably express a PE2-P2A-mCherry construct (HCT116-PE2 cells). We cloned the LDLR137-219 library into an all-in-one lentiviral vector expressing IN-PE2, an improved version of PE2^28^, and infected this library in HCT116-PE2 cells in four biological replicates. We employed an established fluorescent LDL-C uptake screening approach in which four populations are flow cytometrically sorted from each replicate based on LDL-C uptake (lowest 20%, next lowest 20%, highest 20%, next highest 20%) along with unsorted cells as a control (**Figure 1B**)^11,43^. Sorted and unsorted populations were then processed separately to perform NGS preparation of the endogenous LDLR137-219 genomic region and pegRNA-reporter construct.

### Activity-normalized prime editing screening can accurately measure LDLR variant effects

Our data indicate that activity normalization is an effective approach to account for variable editing efficiency. In unsorted cells, a mean of 6.0% of endogenous alleles showed prime editing (**Figure 1C, Supplemental Table 1**). Editing was highly variable—there was a 17-fold difference in the installation efficiency of variants in the 25^th^ vs. 75^th^ percentile in unsorted cells (**Figure S2**). To determine whether variable endogenous editing was determined more by variable pegRNA representation or editing efficiency of pegRNAs, we analyzed the pegRNA-reporter NGS data. Reporters showed overall mean editing of 10.7% (**Figure 1C, Supplemental Table 1**), while showing high variance with 30% of pegRNAs showing no reporter editing (**Figure S2B**). There was modest correlation between the mean abundance of each pegRNA and mean endogenous editing efficiency in unsorted samples (Spearman r = 0.42, Pearson R = 0.35, **Figure 1D**), suggesting that variable pegRNA representation only partially explains endogenous editing variability. We then calculated normalized pegRNA activity as [(mean pegRNA abundance)*(mean reporter editing efficiency)], a metric that normalizes for the activity of each pegRNA. Normalized pegRNA activity showed substantially higher correlation with endogenous editing efficiency than pegRNA abundance (Spearman r = 0.62, Pearson R = 0.51, **Figure 1E**). Altogether, we conclude that pegRNA activity normalization through a pegRNA-reporter improves imputation of endogenous editing efficiency in a high-throughput library setting.

We then used the BEAN pipeline^11^ to analyze variant effects on LDL-C uptake using the flow cytometrically sorted populations from the LDLR137-219 screen. We ran BEAN on both the endogenous variant abundances, which do not require activity normalization, as well as the pegRNA abundances using reporter editing as activity normalization (**Supplemental Table 1)**. We note that, while BEAN is also capable of accounting for multiple distinct genotypic outcomes of each gRNA, because the vast majority of prime editing reporter outcomes are either unedited or edited to the designed variant^25^, we only considered the frequencies of these two outcomes. Stop-gain variants showed significantly lower BEAN Z-scores than synonymous variants using both endogenous and pegRNA BEAN scores, with missense variants showing intermediate scores (p<0.0001, **Figure 2A, Figure S2C**), indicating robust screen performance on positive and negative controls. While the endogenous and pegRNA Z-scores for all variants showed only moderate concordance (Spearman r = 0.33, Pearson R = 0.51, **Figure 2B**), among the 159 variants found to significantly alter LDL-C uptake using endogenous editing (p<0.05), the correlation between BEAN scores was strong (Spearman r = 0.80, Pearson R = 0.73, **Figure S2D**). We conclude that activity normalized pegRNA data is sufficient to accurately identify variants that robustly alter LDL-C uptake.

**Figure 2.**
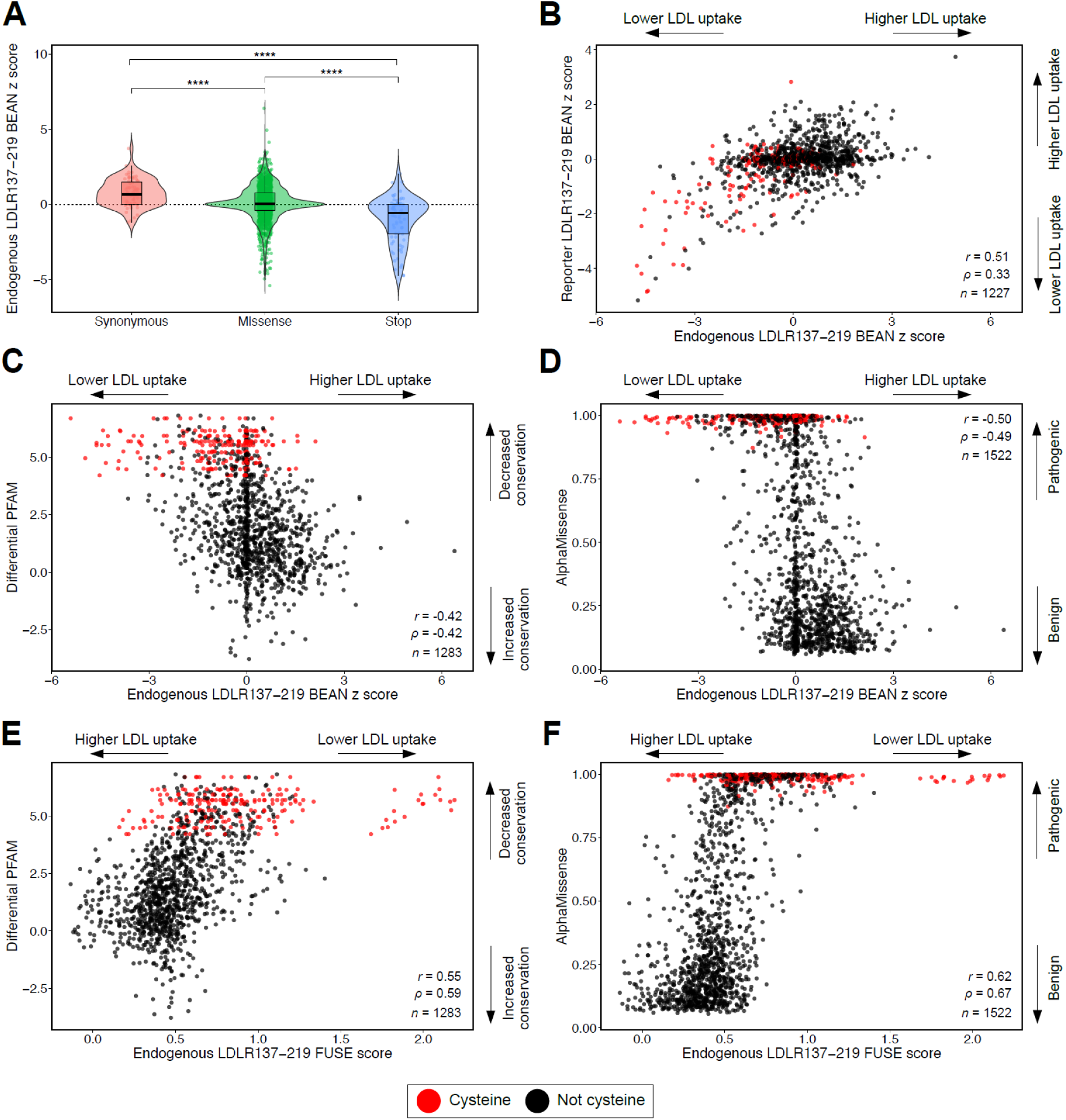
LDLR137-219 screen functional scores reflect domain conservation and computational pathogenicity scores. **A**, Violin plot of BEAN z scores of synonymous variants versus missense (*P* = 2.8 x 10^-8^), missense versus stop-gain (*P* = 8.9 x 10^-9^), and synonymous versus stop-gain (*P* = 5.7 x 10^-15^). Significance was determined by the two-sided Welch *t*-test. **B**, Correlation between LDLR137-219 reporter and endogenous BEAN *z* scores. **C-F**, Scatterplots of **C, E**, differential PFAM scores and **D, F**, AlphaMissense scores versus endogenous LDLR137-219 BEAN *z*-scores (**C-D**) and FUSE (**E-F**) scores. \**P* ≤ 0.05, \*\**P* ≤ 0.01, \*\*\**P* ≤ 0.001, \*\*\*\**P* ≤ 0.0001; n.s., not significant (*P* > 0.05). *r*, *ρ*, Pearson, Spearman correlation coefficient.

The LDLR137-219 region spans highly conserved LDLR class A repeats, and thus we asked to what degree the functional effects measured in our screen reflect disruption of the conserved domain structure. We calculated a conservation score using the PFAM alignment of 244,445 LDLR class A domains. The differential PFAM score for a given variant is the differential log frequency of the variant residue *vs.* the wild-type residue at that position in the PFAM alignment^45^. Aligning the portions of LDLR class A domains 4 and 5 included in the LDLR137-219 library with this PFAM domain architecture, we found robust concordance between variant effects on function and on conservation (Spearman r = -0.42, Pearson R = -0.42, **Figure 2C**), indicating that there is a quantitative relationship between conservation of the domain in general and the contribution of these specific LDLR domains to LDL-C uptake. Based on this finding, we asked to what extent variant effects are captured by the state-of-the-art computational variant pathogenicity prediction model, AlphaMissense. We found that BEAN scores showed strong correlation with AlphaMissense scores (Spearman r = -0.49, Pearson R = -0.50, **Figure 2D**), suggesting that, at least in this highly conserved region of LDLR, prime editing screening and computational modeling converge on a common set of deleterious variants.

While phenotypic measurements from screening assays generally align with clinical outcomes, experimental noise may affect the accuracy of individual variant estimates. We asked whether we could leverage multiple related missense measurements at each position to mitigate this experimental and statistical noise. The FUSE pipeline^31^ estimates the functional impact of any variant at an amino acid residue using shrinkage estimation to reduce noise and then leverages a substitution matrix of missense effects across 115 published deep mutational scans to adjust observed scores of missense variants within a given starting position. FUSE-adjusted BEAN scores from the LDLR137-219 screen showed improved correlation with differential PFAM scores (Spearman r = 0.59, Pearson R = 0.55, **Figure 2E, Supplemental Table 1**) and AlphaMissense scores (Spearman r = 0.67, Pearson R = 0.62, **Figure 2F**) as compared to the equivalent BEAN scores, suggesting that FUSE effectively reduces noise in our dense functional screening data. Altogether, the LDLR137-219 data demonstrates that activity-normalized prime editing screening is capable of accurately scoring variant effects from high-throughput screens.

### Full LDLR activity-normalized prime editing screening

We next extended our screen to assess variant effects across the full length of the *LDLR* coding sequence. Several technical aspects were altered in the design and implementation of LDLR full length (LDLR-FL) library screening. Because sequencing endogenous edits across the length of *LDLR* is infeasible, we focused data collection on pegRNA-reporter pairs and only included pairs in which editing efficiency readout was possible within the constraints of a 300-nt oligo library. To minimize pegRNA number while ensuring high-coverage screening, we selected four missense variants per position predicted to provide highest imputation accuracy using the GLIDE approach (manuscript in preparation). In addition, we included every missense variant present in ClinVar and in ∼470,000 individuals with whole exome sequences in the UK Biobank, and 600 synonymous and stop-gain variants distributed across the coding region along with 305 non-targeting negative controls. We used PRIDICT2.0^30^ to determine optimal pegRNAs for each edit. Our final library contained 6,000 total pegRNAs including 4,495 missense variants spanning 816 of the 860 *LDLR* amino acids (**Supplemental Table 2**).

We cloned the LDLR-FL library into an IN-PE2 vector modified to append the pegRNA-stabilizing La(1-194) protein to the prime editor (IN-PE2-SSB), an approach shown to improve prime editing efficiency^27^. We performed four biological replicates of fluorescent LDL-C uptake screening in HCT116 cells engineered with knockout of the lentivirus-silencing TASOR gene^46^ and stably expressing PE2 using the same sorting paradigm as our pilot screen and preparing NGS libraries of pegRNAs and paired reporters.

The LDLR-FL library provided robust variant editing. Read counts across sorted populations within a replicate and between biological replicates showed high correlation (Spearman r within a replicate 0.79-0.89, between replicates 0.69-0.79, **Figure S3A**). Reporter editing was robust but highly variable. Median editing was 53%, with 1,297 pegRNAs (21.6%) showing <10% editing and 976 pegRNAs (16.3%) showing >90% editing (**Figure 3A, Supplemental Table 2**). Although pegRNAs were pre-selected for high PRIDICT2.0 scores, reporter editing still showed moderate correlation with PRIDICT2.0 score (Spearman r = 0.35, Pearson R = 0.36, **Figure 3B**). Among pegRNA features, the RT template length had a robust negative correlation with the length of the RT template (Spearman r = - 0.39, Pearson R = -0.38, **Figure 3C**). Longer RT templates are required when variants are distal to the PAM, revealing that the random distribution of PAM sequences is a major driver of variable prime editing efficiency.

**Figure 3.**
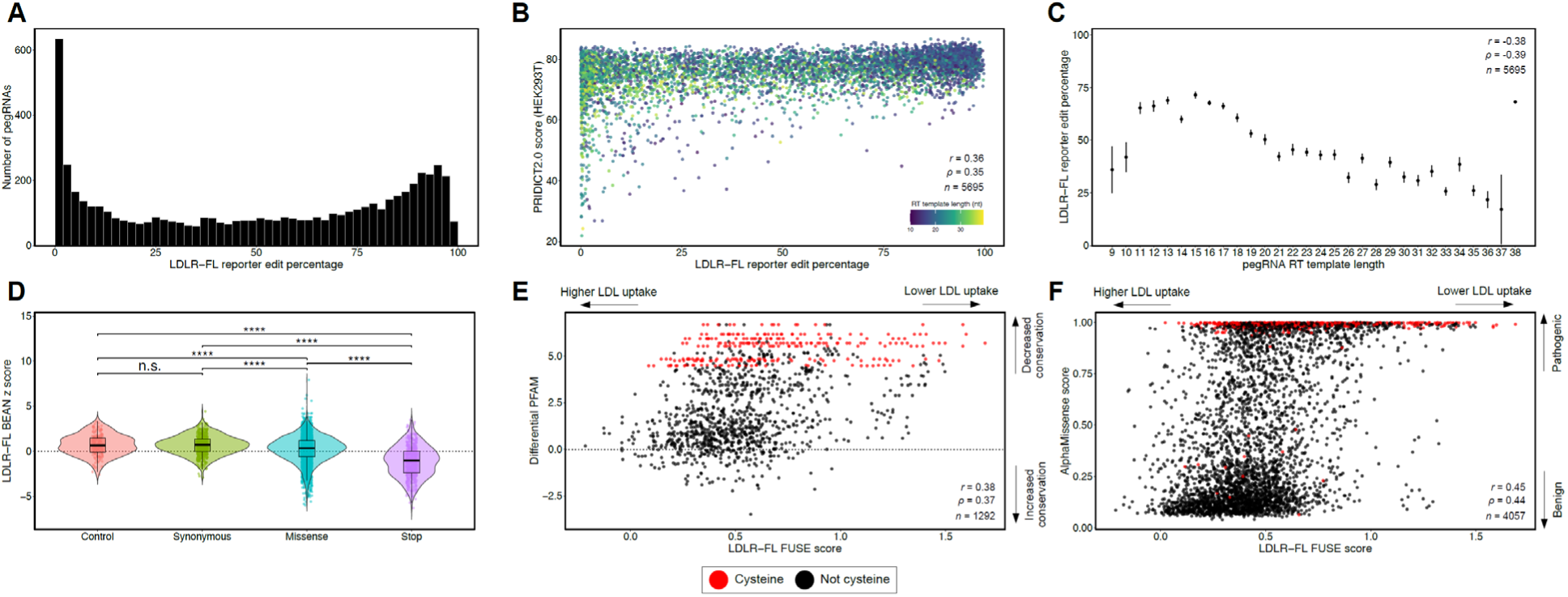
Analysis of correlates of prime editing efficiency and functional scores from LDLR-FL screen. **A**, Histogram of LDLR-FL pegRNA reporter edit fractions as determined through BEAN analysis of replicates. **B**, Scatterplot of PRIDICT2.0 scores versus LDLR-FL reporter edit fraction. **C**, LDLR-FL reporter edit fraction versus pegRNA RT template length. **D**, Violin plot of BEAN z scores of control versus synonymous variants (*P* = 0.83), synonymous versus missense variants (*P* = 7.0 x 10^-16^), and missense versus stop-gains (*P* < 2 x 10^-16^). Significance was determined by the two-sided Welch *t*-test. **E-F**, Scatterplot of **E**, differential PFAM and **F**, AlphaMissense versus LDLR-FL FUSE scores. \**P* ≤ 0.05, \*\**P* ≤ 0.01, \*\*\**P* ≤ 0.001, \*\*\*\**P* ≤ 0.0001; n.s., not significant (*P* > 0.05). *r*, *ρ*, Pearson, Spearman correlation coefficient.

We then processed these counts using BEAN and FUSE on the LDLR-FL dataset (**Supplemental Table 2**). Stop-gain and missense variants showed significantly lower Z scores than control and synonymous variants (**Figure 3D**), suggesting, as expected, that these variants disrupt LDLR function. Variants shared between the LDLR137-219 and LDLR-FL libraries showed robustly correlated BEAN and FUSE scores (**Figure S3B-C**). BEAN and FUSE scores for missense variants also correlated robustly with differential PFAM scores within LDLR class A repeats and AlphaMissense scores (**Figure 3E-F, Figure S3D-E**). In both cases, FUSE scores showed stronger correlation than BEAN scores, suggesting that FUSE successfully denoised the prime editing data using the multiple missense variants at each position.

### Calibration and Impact of Functional Evidence on LDLR Variant Classification

We next asked whether LDLR-FL functional scores could be used to distinguish pathogenic *LDLR* missense variants. ClinVar pathogenic and likely pathogenic missense variants had significantly higher functional scores as compared to benign and likely benign variants (p<0.0001), **Figure 4A**), with variants of uncertain significance (VUS) and those with conflicting annotations having intermediate scores. We also compared FUSE scores with mean LDL-C levels in participants with rare *LDLR* missense variants in the UK Biobank cohort. FUSE scores for variants with mean LDL-C >190 mg/dL were significantly higher than for those <190 mg/dL and showed moderate overall correlation (p<0.0001, **Figure 4B, Figure S4**). FUSE scores for variants within participants diagnosed with CAD were higher than for those without CAD, although this difference was not statistically significant (p=0.1, **Figure 4C**), possibly due to the relative sparsity and multi-factorial etiology of CAD as compared to LDL-C levels. Thus, FUSE scores are associated with LDLR variant pathogenicity. Importantly, the LDLR-FL screen directly measured the function of 381 of the 452 (84.3%) LDLR missense variants present in the UKB cohort, while our earlier base editing screens only measured 55 (12%, **Figure S4**), showcasing the superior coverage of prime editing screens.

**Figure 4.**
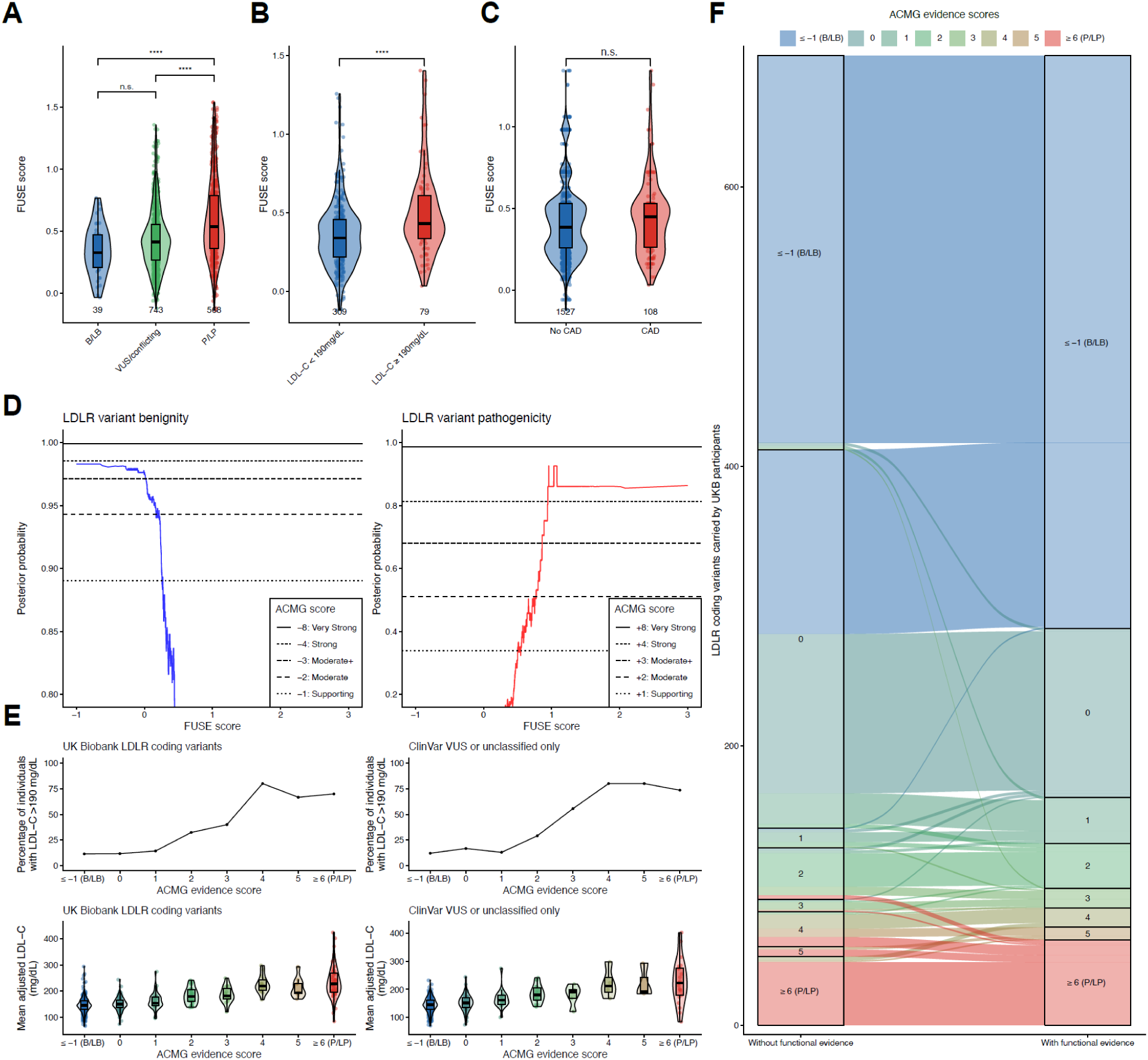
LDLR prime editing functional scores enhance clinical variant interpretation. **A,** Violin plots of FUSE-scored variants stratified by ClinVar classifications (*P* = 0.053), (*P* = 1.1 x 10^-6^), (*P* = <2.0 x 10^-16^). **B,** Violin plots of FUSE-scored variants found in UK BioBank participants with LDL-C <190 mg/dL versus ≥190 mg/dL (P = 1.0 x 10^-5^). **C,** Violin plots of FUSE-scored variants found in UK BioBank participants with or without a diagnosis of coronary artery disease (CAD) (*P* = 0.1). **D,** Posterior probability scores for LDLR variant benignity and pathogenicity versus LDLR-FL FUSE scores. **E,** Distribution of LDL-C levels for UK Biobank LDLR coding variants and ClinVar VUSs as a function of ACMG/AMP evidence scores. **F,** Alluvial diagram illustrating ACMG/AMP evidence category assignments before and after incorporating functional evidence from the LDLR prime-editing screen.

We then sought to understand the impact of our LDLR-FL functional scores on clinical variant classification using the ACMG/AMP sequence-variant-interpretation (SVI) framework^37^. To enable quantitative incorporation of experimental functional data into variant classification, we first calibrated the predictive strength of functional evidence derived from the LDLR-FL screen using a recently described framework^36^. Calibration of LDLR-FL FUSE scores against ClinVar reference variants yielded posterior probability curves that closely followed the expected monotonic relationship between assay scores and the likelihood of pathogenicity (**Figure 4D**). The estimated posterior probabilities provided clear functional score thresholds corresponding to Supporting, Moderate, and Strong levels of evidence for variant classification within the ACMG/AMP SVI. This result demonstrates that the functional data is sufficiently concordant with existing clinical classifications to be directly translated into standardized evidence strengths for clinical interpretation.

We next measured how many *LDLR* variants which were uncertain or absent from ClinVar might have sufficient evidence to be classified as pathogenic or benign. We integrated four forms of evidence which we could electronically assemble without a manual curated review of clinical variant summaries, including computational scores (AlphaMissense)^38^, population enrichment of disease at the variant level^39^, clinical context from previously classified variants within the same amino acid residue^47^, and the functional evidence developed from our LDLR-FL screen. We combined these forms of evidence using a Bayesian adaptation of the ACMG/AMP SVI framework^37^ as described in the **Methods**, noting that a comprehensive evaluation of all available forms of evidence would be required for a final diagnostic variant classification. Applying the calibrated functional evidence to *LDLR* variants observed in the UK Biobank revealed strong concordance between the ACMG functional evidence scores and clinical lipid phenotypes (**Figure 4E**). Across all coding variants, mean LDL-C levels of carriers increased progressively with higher ACMG evidence scores, accompanied by a corresponding rise in the proportion of individuals with LDL-C ≥ 190 mg/dL. Notably, among the rare *LDLR* variants within the UK Biobank currently labeled as VUS, conflicting, or unclassified in ClinVar, 322 (74.2%) had evidence sufficient for reclassification after integrating all four evidence types: 301 (69.3%) to benign/likely benign (B/LB) and 21 (4.8%) to pathogenic/likely pathogenic (P/LP) after integration of all four evidence types (**Figure 4E, Supplemental Table 3**), underscoring the potential of calibrated functional data to resolve clinical uncertainty. Altogether, these relationships validate the biological and clinical relevance of the composite ACMG scoring system that integrates population, computational, clinical, and functional evidence.

We further examined the effect of adding functional evidence on overall ACMG classifications. Incorporation of the calibrated LDLR-FL FUSE scores altered the ACMG evidence assignments for 210 of 694 (30.3%) rare *LDLR* variants observed in UK Biobank (**Figure 4F, Supplemental Table 4**). Among these, 133 variants (19.1%) gained sufficient evidence to meet the threshold for B/LB classification, while 16 variants (2.3%) were upgraded to P/LP. These shifts highlight the substantial impact of well-calibrated experimental functional data in improving the resolution and accuracy of variant classification, particularly for variants previously categorized as having uncertain significance.

### LDLR class A repeat 5 is enriched in gain-of-function variants

We next analyzed missense variant effects by domain. LDLR class A repeats 3-5 (LDLRA3-5) showed the highest mean LDLR-FL FUSE scores, while C-terminal regions including residues known to mediate interaction with the LDLR-degrading E3 ligase MYLIP^48^ and the EGF-like repeat 3 and clustered O-linked oligosaccharide domains showed the lowest (**Figure 5A**). LDLRA3-5 also showed the highest density of variants that significantly decreased LDL-C uptake (**Figure 5B**). These trends were reinforced when examining variants individually (**Figure 5C**). Of the 25 most deleterious variants, 14 altered highly conserved cysteines or aspartic acids within LDLR class A repeats. Because the conserved cysteines form disulfide bridges within each LDLR class A repeat, conferring structural stability, and the aspartic acid residues mediate LDLR-apolipoprotein electrostatic interactions, this result matches our expectations. More surprisingly, the four strongest gain-of-function (GOF) variants in the entire gene all resided within LDLRA5 (**Figure 5C**).

**Figure 5.**
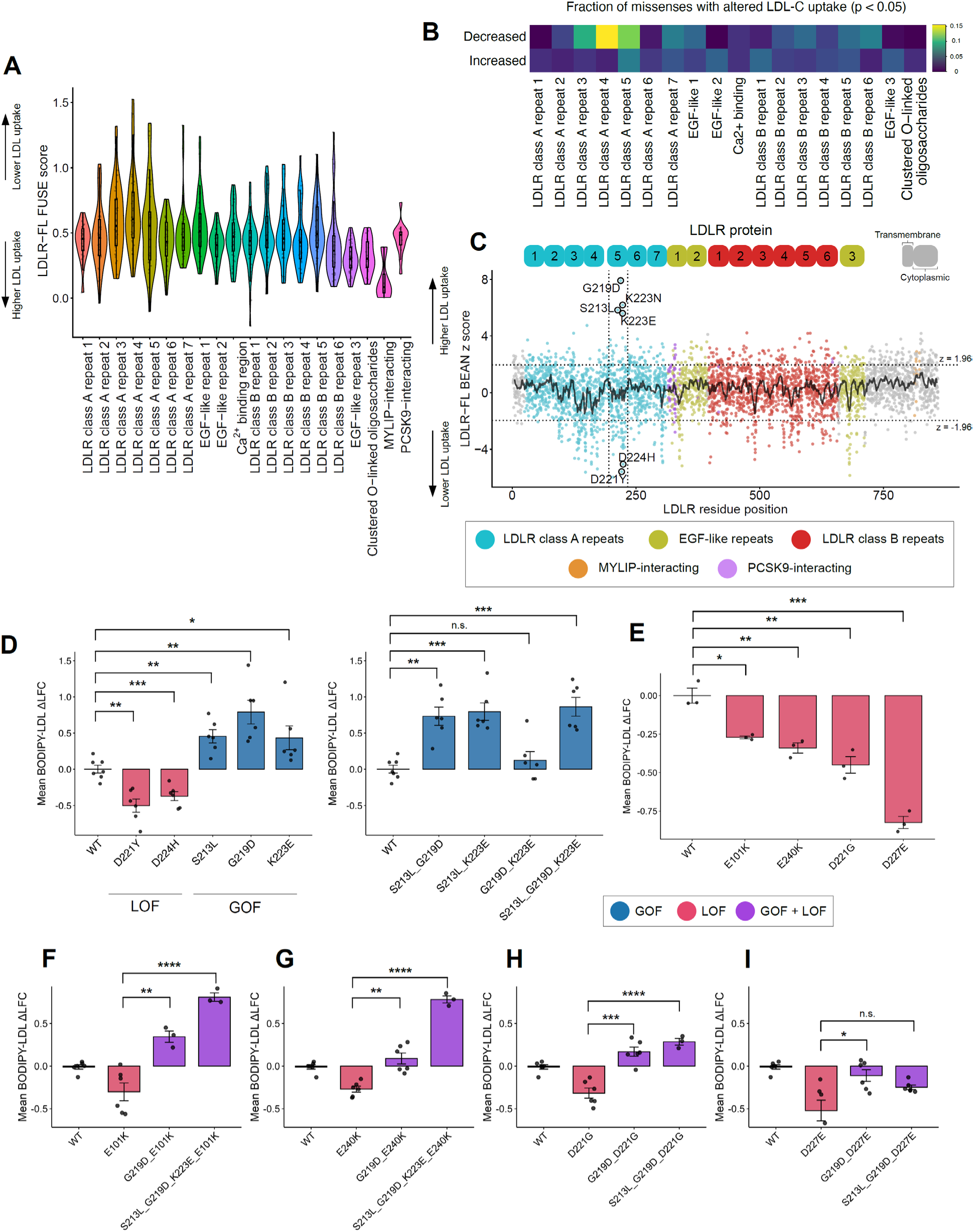
LDLR class A repeat 5 harbors LDL uptake-increasing variants that functionally complement pathogenic variants. **A**, Violin plot of LDLR-FL FUSE scores for annotated LDLR structural domains. **B**, Heatmap of missense variant fractions with significant LDLR-FL BEAN z scores (p<0.05) for annotated LDLR structural domains. **C**, Scatterplot of LDLR-FL BEAN *z*-scores colored by structural domain. Variants chosen for individual functional validation are circled in black. **D-I**, Boxplot of BODIPY-LDL ΔLFC for selected LDLR loss-of-function (LOF), gain-of-function (GOF) variants and GOF variant combinations (**D**), ClinVar pathogenic LDLR loss-of-function (LOF) variants (**E**), and LOF-GOF variant combinations **(F-I)**. \**P* ≤ 0.05, \*\**P* ≤ 0.01, \*\*\**P* ≤ 0.001, \*\*\*\**P* ≤ 0.0001; n.s., not significant (*P* > 0.05).

To investigate these strong gain-of-function (GOF) variants further, we cloned three candidate GOF variants (S213L, G219D, and K223E) and two adjacent deleterious variants (D221Y, D224H) into an inducible lentiviral LDLR-mCherry fusion vector. All three GOF variants significantly increased LDL-C uptake over LDLR-wt, while the deleterious variants decreased LDL-C uptake (**Figure 5D**). We also tested all combinations of the GOF variants, finding that three of four combinations significantly increased LDL-C uptake with the triple mutant LDLR^S213L,G219D,K223E^ showing strongest uptake (**Figure 5D**). We note that while LDLR^S213L,G219D,K223E^ showed stronger uptake than any individual GOF variant, the relative increase was sub-additive, suggesting some functional redundancy among these variants.

We subsequently tested whether GOF variants could restore LDLR function when concurrently installed with pathogenic LOF variants. We selected four LOF variants that: (1) are classified as pathogenic or likely pathogenic in ClinVar; (2) show elevated mean LDL-C in ≥5 heterozygous carriers in the UK Biobank; (3) reduced LDL uptake in our screen (FUSE >0.4); and (4) are predicted to be damaging by AlphaMissense (>0.4). When expressed individually, these LOF variants decreased LDL uptake in HepG2 cells compared to wild-type (**Figure 5E**). We then generated combinatorial variant constructs containing individual LOF variants as well as either G219D or a combination of GOF variants. For all four LOF variants, *cis* complementation with GOF variants significantly improved LDL uptake, often to levels exceeding wild-type expression, highlighting their potential to functionally compensate for pathogenic mutations (**Figure 5F-I**).

### LDLRA5 GOF variants are predicted to increase intermolecular contacts with APOB

We next investigated possible mechanisms driving these GOF variants. To evaluate effects on LDLR stability, we modeled mutations using the X-ray structure of the LDLR extracellular domain^49^ with FoldX^50^ and DDMut^51^. To assess effects on LDLR-APOB binding, we used a cryo-EM structure of the LDLR-class A repeats bound to the apolipoprotein B (APOB) beta-belt^49^ and predicted affinity changes with DDMut-PPI^52^, as well as interatomic contact changes using the webserver Arpeggio ^53^ (**Figure 6A**).

**Figure 6.**
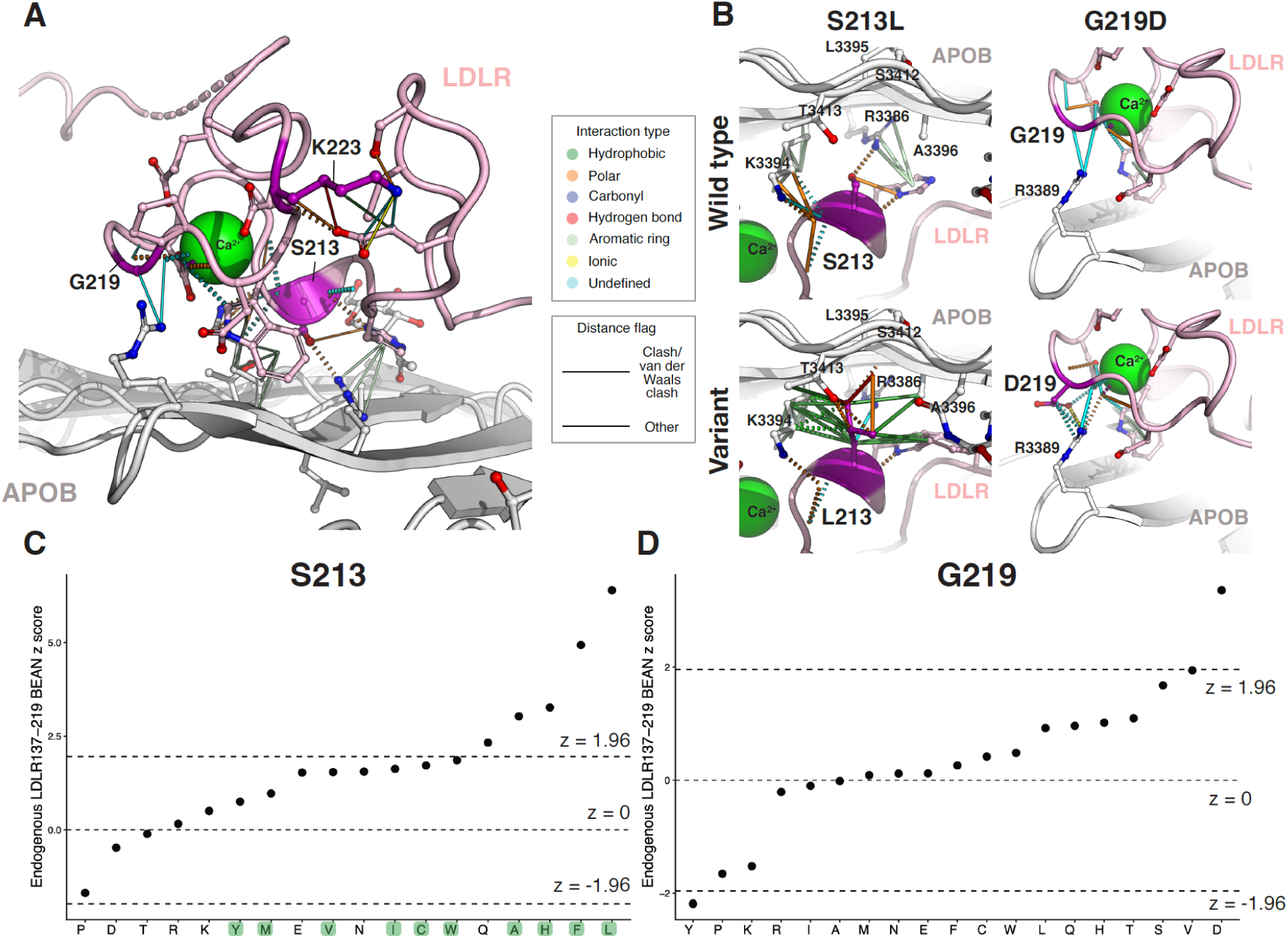
Strong LDLR GOF variants generate novel LDLR-APOB inter-atom contacts. **A**, LDLR-APOB cryo-EM co-structure with LDLR residues of interest highlighted in purple. **B**, Local interatomic interactions in wild-type and variant structures for GOF variants S213L and G219D. Atoms are colored by elements (O, red; N, blue; S, yellow) and calcium heteroatoms are colored light green. **C-D,** Scatterplots of endogenous LDLR137-219 BEAN Z-scores for variants introduced at S213 and G219 respectively. Hydrophobic residues are highlighted in green.

Although none of DDMut, DDMut-PPI, or FoldX predicted increased LDLR stability or LDLR-APOB affinity for GOF variants LDLR^S213L^, LDLR^G219D^, and LDLR^K223E^ (**Figure S5**), Arpeggio revealed that LDLR^S213L^ and LDLR^G219D^ generate novel interatom contacts at the LDLR-APOB interface. We find that LDLR^S213L^ generates twelve additional predicted hydrophobic interactions and LDLR^G219D^ generates a novel ionic interaction between LDLR^D219^ and APOB^R3389^ (**Figure 6B**). LDLR^K223E^ lies more than 5 Å from APOB, and Arpeggio does not predict any intermolecular interactions between LDLR^K223E^ and APOB, suggesting that LDLR^K223E^ may enhance LDL uptake through a mechanism independent of LDLR-APOB affinity.

To gain further insight into the attributes contributing to GOF variants, we examined BEAN Z scores for all 19 missenses for S213 and G219, as these residues were included in the LDLR137-219 library. Four of the five S213 missenses that induce significant GOF convert serine to a hydrophobic residue (leucine, phenylalanine, histidine, and alanine, **Figure 6C**), supporting the idea that increased hydrophobic interactions play a role in the improved function at this position. On the other hand, G219D is the only significant GOF swap at position 219 (**Figure 6D**), and the other negative charge-gain variant, G219E, has neutral effect. Taken together, these observations suggest that negative charge as well as other factors such as steric effects contribute to the unique function-enhancing properties of G219D.

We then experimentally probed LDLR GOF variant mechanisms. First, we used flow cytometry of variant LDLR-mCherry-transduced cells to evaluate LDLR variant stability. While the LOF LDLR^D221Y^ variant showed significantly lower mCherry fluorescence, suggesting impaired stability, none of the three top GOF variants showed significantly increased stability (**Figure S6A**). Because the primary role of LDLR class A repeats is in binding to the LDL particle, we evaluated whether increased uptake resulted from enhanced LDL association. We transduced K562 cells with GOF or LOF variants and evaluated fluorescent LDL association in the presence of Pitstop 2, a potent clathrin-mediated endocytosis inhibitor^54^, which we confirmed prevented LDL uptake (**Figure S6B**). We found that LDLR^G219D^ and LDLR^K223E^ increased and LDLR^D221Y^ and LDLR^D224H^ decreased LDL association (**Figure S6C**). Altogether, structural and functional evidence converge to suggest that LDLR GOF variants may act to increase association between LDLR and LDL particles through enhancing affinity with APOB.

### Comparison of approaches to measure LDLR variant effects

Two recently published manuscripts used base editing and cDNA approaches to functionally screen *LDLR* variants at scale. We compared functional scores for 620 (Ryu *et al.*^11^ base editing), 15,507 (Tabet *et al.*^12^ cDNA), and 5,186 (combined output of LDLR137-219 and LDLR-FL prime editing screens) *LDLR* missense variants with AlphaMissense scores and UKB carrier LDL-C levels (**Figure 7A-B, Supplemental Table 5**). Scores from the three functional screening approaches show moderate concordance (Spearman r = 0.30-0.67, Pearson R = 0.39-0.65). Scores from Ryu *et al.* associated strongly with UKB carrier LDL-C levels and AlphaMissense scores, with prime editing and Tabet *et al.* scores showing robust but slightly weaker associations. We note that Tabet *et al.* scored 446 of 452 UKB missense variants compared to 381 and 55 variants scored by the combined prime editing libraries and Ryu *et al.* respectively (**Figure S4B**).

**Figure 7.**
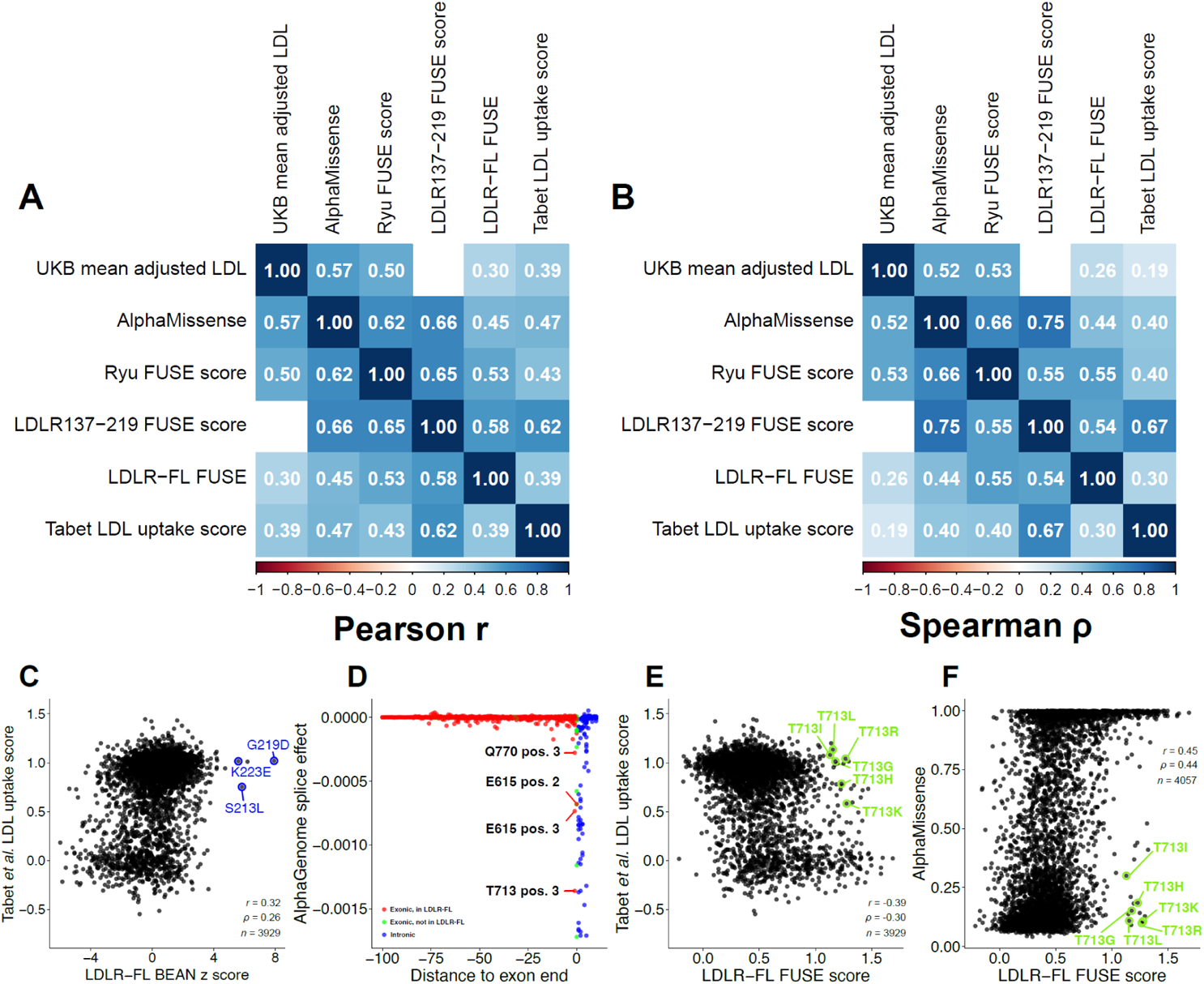
Comparison of LDLR variant effect data from CRISPR screening and cDNA deep mutational scanning. **A-B,** Pearson **(A)** and Spearman **(B)** correlations among LDLR functional scores, AlphaMissense predictions, and UKB LDL-C levels. **C,** Correlation plot comparing LDLR-FL BEAN Z-scores with Tabet *et al.* LDL uptake functional scores. Gain-of-function variants from the LDLR-FL screen are circled in blue. **D,** AlphaGenome predicted splice effects of splice donor variants with strongest exonic splice donor variants labeled. **E-F,** Scatterplots of LDLR-FL FUSE scores vs. Tabet *et al.* (**E**) and AlphaMissense (**F**) scores with all variants at the top splice donor site (T713) circled in green. *r*, *ρ*, Pearson, Spearman correlation coefficient.

In spite of the overall similarity, the prime editing data differed from the Tabet *et al*. data in several notable ways. First, the strong GOF residues identified in the LDLR-FL screen failed to stand out as hypermorphic in the Tabet *et al.* data (**Figure 7C**). It is unclear what underlies this discrepancy, but Tabet et al. used a single flow cytometry gate of top 20% of cells, which might hinder sensitivity to identify GOF residues.

Second, candidate splice-altering residues showed distinct activity. We used AlphaGenome^55^ to predict the effect on splice-site usage of every possible single-nucleotide change in the *LDLR* coding sequence and proximal intronic regions. While the vast majority of the strongest predicted splice-altering variants are in intronic splice donor and acceptor regions (**Figure 7D**, **Figure S7A**), the third nucleotide in T713, which lies two nucleotides away from the exon 14 splice donor, was predicted to abrogate splicing with similar strength to splice donor disruptions (**Figure 7D**). Accordingly, all six T713 missenses in the LDLR-FL library showed strong loss-of-function phenotypes, while these variants appeared neutral in the Tabet *et al.* dataset and received low AlphaMissense scores (**Figure 7E-F**). Because neither cDNA-based screening nor AlphaMissense incorporate splicing information, these results suggest that our prime editing strategy is better suited to detecting functional consequences of splice-altering coding variants. AlphaGenome predicted weaker splice-altering effects in two additional residues, E615 and Q770. Prime editing showed slightly more deleterious effects of Q770 missenses than Tabet *et al.* and AlphaMissense (**Figure S7B-C**), although the discrepancies were more modest than those observed at T713. Further experimentation is required to validate that observed differences result from splice alteration. Altogether, these comparisons reveal that activity-normalized prime editing data provides equivalent quality data to cDNA-based deep mutation scanning with a less laborious workflow and advantages in detecting candidate splice-altering variants.

## Discussion

In this work, we introduce a framework to perform dense functional analysis of coding variant effects while accounting for the variable installation efficiency inherent to PE. Activity normalization is valuable in PE screening for two reasons. First, PE shows highly variable installation efficiency, and this variability is only partially accounted for by state-of-the-art pegRNA efficiency prediction^30^, necessitating an empirical solution to estimating installation efficiency. Prior screens using integrated reporter constructs have filtered low-editing pegRNAs^14^, but accounting for the entire spectrum of editing frequency improves imputation accuracy. Second, improved prime editors are being introduced at breakneck pace. We incorporated several advances reported in the intervening period between designing LDLR137-219 and LDLR-FL screens and observed substantially improved median editing efficiency with the updated editor architecture. Since performing the LDLR-FL screen, several additional improvements have been described^56,57^. Being able to rapidly deploy new PE architectures that may show distinct activity patterns while measuring and accounting for variable efficiency in the same experiment is valuable.

We also introduce a pipeline to reduce the experimental burden of variant scanning through optimizing the selection of missense variants (GLIDE) and improving the estimation of functional effects by collectively analyzing these measurements (FUSE). Full saturation mutagenesis of LDLR would require greater than 18,000 variant effect measurements. It is difficult to obtain high-quality flow cytometric screening data for such a large number of variants, especially as incomplete prime editing efficiency necessitates averaging phenotypic data from a large pool of cells per pegRNA. Our results demonstrate that FUSE-processed missense scores correlate more strongly with computational and clinical metrics of LDLR variant function than BEAN scores, emphasizing the value of this approach. It is also worth noting that PE screening is better suited to rational library design than base editing screening, which can only install 1-2 variants per codon^11^.

Finding the optimal balance of installation *vs.* imputation is a parameter we did not address in this study. This balance depends on assay-specific parameters such as throughput and noise. Cell survival-based screens allow higher throughput, flow cytometric screens have intermediate throughput, and high-content phenotypes such as single-cell RNA-seq and imaging show lowest throughput and thus may benefit even more from rational variant selection. The balance of installation *vs.* imputation also depends on the goals of the screen. One focal point of this work was the identification of gain-of-function variants in LDLRA5. Because GOF variants rely on the addition of specific new chemical properties, they present more of a challenge to imputation approaches than loss-of-function variants^58^. For example, G219D is the only significant GOF variant identified at position 219, and we present structural modeling suggesting that a gained ionic interaction with APOB^R3389^ at the interaction interface may underlie the unique phenotypic properties of the aspartic acid. However, the FUSE score of G219D is the 11^th^ lowest of G219 missenses, presumably because FUSE is trained to identify typical missense relationships averaged across thousands of residues, and thus it is poorly suited to identifying unique effects that depend on a specific protein-protein interface. Similarly, AlphaMissense scores G219D as the 6^th^ least pathogenic missense variant at its position, possibly because the focus of AlphaMissense training is in predicting pathogenic variants and not in flagging GOFs^59^. Even structural prediction approaches such as DDMut, DDMut-PPI, and FoldX fail to correlate with our screen data in the identification of strong GOF variants. Altogether, due to the rarity of GOF variants combined with the lack of adequate computational tools to impute them, screens to identify GOF variants should be more exhaustive than those focused exclusively on scoring pathogenicity of LOF variants.

It is striking that the four strongest GOF variants all reside within LDLRA5. A prior study used a candidate approach to produce *LDLR* variants that escape degradation by MYLIP and PCSK9 and show enhanced efficacy at reducing serum LDL-C levels upon adeno-associated virus infection of hypercholesterolemic mice^60^. Our study supports the finding that variants in MYLIP-interacting residues enhance LDL-C uptake, although these variants show weaker effects than the strongest LDLRA5 GOF variants. On the other hand, variants in PCSK9-interacting residues do not increase LDL-C uptake in our screen. HCT116 express PCSK9 at low levels, so it is possible that our model system underestimates the effects of PCSK9 and thus is unable to properly evaluate candidate GOF variants at PCSK9-interacting residues.

Within LDLR class A repeats, there is a plausible explanation of the salience of LDLRA5. LDLR primarily engages with two ligands on LDL particles, APOB and APOE. Every LDL particle contains APOB, while <10% of LDL particles typically carry APOE^61^. However, APOE has a 20-fold higher affinity for LDLR than APOB^62^. Previous work has identified LDLRA3-7 as essential for binding APOB^49,63^, and LDLRA5 is the only structural element shown to be essential for APOE binding^64^. So, one hypothesis for the salience of GOF residues on LDLRA5 is that they are uniquely capable of enhancing interaction with both APOB and APOE-containing LDL particles.

The LDLR-APOB co-structure reveals that the strongest GOF variants sit at the binding interface of LDLR and APOB^49^, suggesting that increasing affinity of this interaction and potentially the cognate LDLR-APOE interaction is a plausible mechanism by which such variants enhance LDL uptake. Our structural modeling provides further support for this idea, as both S213L and G219D variants are predicted to add intermolecular interactions between LDLR and APOB. Of particular note, G219D forms an ionic interaction with APOB^R3389^. Ionic interactions between negatively charged residues on LDLR and positively charged APOB residues are known to play a key role in their interaction^49^, and in fact, G219D has previously been shown to enhance the affinity of LDLR-APOE binding^65^. The lack of a solved LDLR-APOE structure prevents computational analysis of LDLR variant-mediated affinity changes with APOE. Furthermore, docking solved structures of LDLR and APOE is unlikely to yield meaningful results because current protein-protein docking methods ignore the lipidated context in which APOE is natively found, which dramatically alters its structure. It will be important to confirm through biochemical assays such as surface plasmon resonance that the GOF variants we identify do indeed act by increasing the affinity of LDLR-APOB/APOE interaction.

Our experiments demonstrate that introducing GOF LDLR variants can compensate for LOF variants known to cause FH, at least in a cellular model. These findings provide preliminary evidence in support of a novel therapeutic approach for FH, the use of genome editing to complement FH variants. This approach shares the same key advantage, one-time dosage, as the recently demonstrated base editing inactivation of PCSK9^66^. It also shares similar drawbacks. We show that this approach is effective at neutralizing four moderate LDLR LOF variants. While we expect this approach to generalize widely to moderate LOF variants, we would expect this approach not to be effective at complementing stop-gain variants or those that severely affect LDLR folding or function. This is similar for PCSK9 inhibition, which acts through upregulating *LDLR* expression and thus cannot overcome defective LDLR. It will be important to evaluate GOF complementation in an *in vivo* setting to compare its LDL-C reduction efficacy with alternative strategies such as PCSK9 inhibition and to rule out immunogenicity of the altered amino acid(s).

In addition to insights into GOF variants, our work, alongside other recent studies^11,12^, shows the potential of functional screens to improve clinical classification of *LDLR* variants. It is striking that, with application of the ACMG/AMP SVI framework, 74.2% of UKB-observed LDLR variants without a definitive ACMG classification meet criteria for classification. While the vast majority of these variants are reclassified as benign, 21 variants (4.8% of those observed in UKB) have evidence that meets the threshold for a pathogenic classification, suggesting that the combination of functional screening with other evidence types has potential to improve diagnosis and prioritize treatment for individuals with FH. It will be fruitful to combine data from this work and the other recent *LDLR* variant scans to build robustness in functional scores to more confidently guide clinical application. We also note that our scores rely on capturing differences in LDL-C uptake in a single cell type under a single environmental condition. As LDLR binds to a variety of apolipoproteins that populate distinct lipoproteins at differing ratios, our screens are unlikely to reflect the full spectrum of physiological effects of LDLR variants *in vivo*.

We also note that *LDLR* variant scores show quantitative association with carrier LDL-C levels beyond what is captured in the binary variant pathogenicity classifications for FH. Our combined prime editing FUSE scores show moderate correlation with adjusted LDL-C levels in UKB carriers (Pearson r = 0.31), as has been found for scores from base editing screening^11^. Aggregated ACMG evidence scores also correlate with carrier LDL-C levels. For example, <10% of LDLR variant carriers with ACMG score of 1 or lower and >75% of carriers with scores of 4 and 5 have adjusted LDL-C at or above the FH threshold of 190 mg/dL, indicating that these scores carry information beyond what meets the threshold of definitive clinical evidence. Not only do these findings suggest that improving evidence of all types could lead to further diagnostic yield, they also suggest that future efforts could utilize information on *LDLR* variants along with additional information such as polygenic risk scores^67^ to provide personalized LDL-C estimates.

Altogether, activity-normalized prime editing offers a powerful approach to dissect the effects of genetic variants on disease. Through deep profiling of *LDLR* variant effects, this work advances genetic diagnosis of FH as well as offering a novel approach to increase LDL-C uptake capacity.

## Acknowledgments

The authors thank Grigoriy Losyev for technical assistance. BioRender schematics were used to generate Figure 1.

## Sources of Funding

Funding for this work was obtained from UM1HG012010 (L.P., R.I.S.), 1R01HL164409 (L.P., C.A.C., R.I.S.), 1R01GM143249 (R.I.S.), 1R56HG012681 (C.A.C., R.I.S.), and the American Heart Association 24TPA1300072 (C.A.C., R.I.S.).

## Disclosures

L.P. has financial interests in Edilytics, Inc., and SeQure Dx, Inc. G.S. is a scientific advisor of Prime Medicine and Co-founder of Nerai Bio. Authors’ interests were reviewed and are managed by MassGeneral Brigham HealthCare in accordance with their conflict of interest policies.

## Supplemental Figures

**Figure S1.**
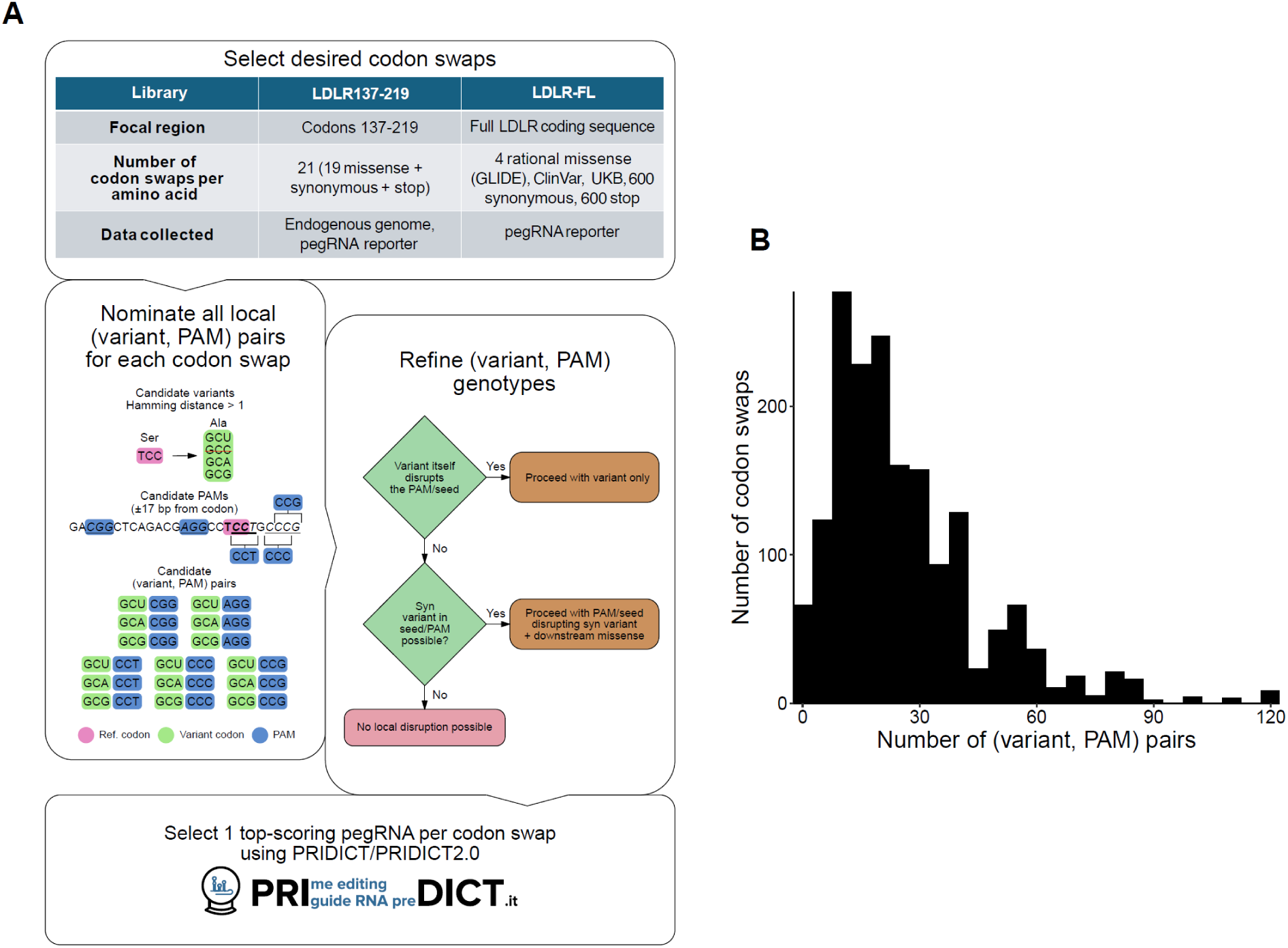
LDLR prime editing library design. **A,** Schematic of the LDLR137-219, LDLR-FL libraries and pegRNA selection workflow. **B,** Histogram of codon swaps by the number of (variant, PAM) pairs.

**Figure S2.**
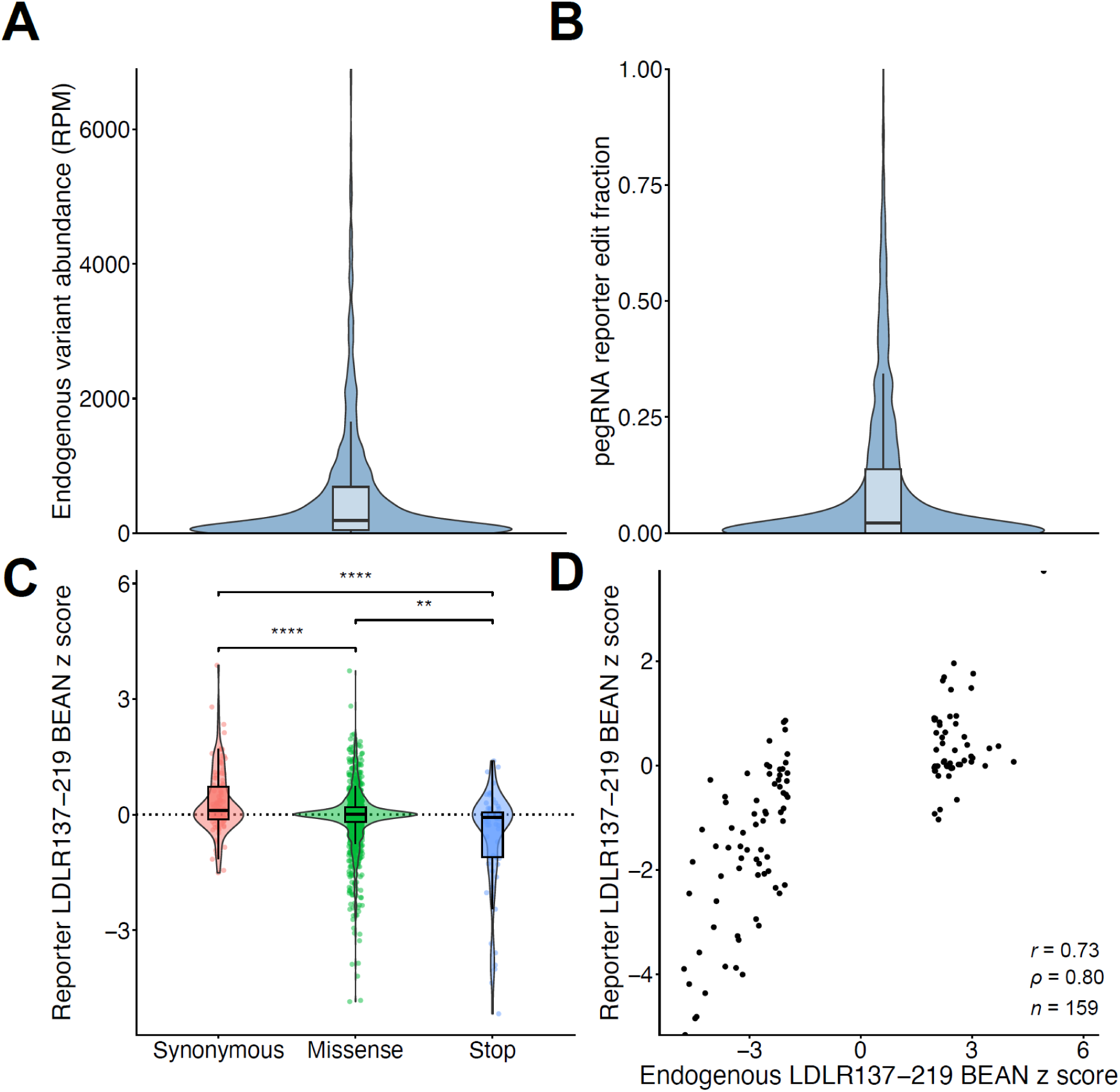
Benchmarking the ANPE pipeline with LDLR137-219. **A-B,** Violin plot for endogenous variant abundance (**A**) and pegRNA reporter edit fraction (**B**) for LDLR137-219. **C,** Violin plot of LDLR137-219 BEAN z-scores of synonymous versus missense (*P* = 2.1 x 10^-5^), missense versus stop (*P* = 0.0025), and synonymous versus stop (*P* = 4.6 x 10^-6^) for the genomic reporter. **D,** Correlations of reporter and endogenous BEAN z-scores only for significant values (|z| < 1.96). *r*, *ρ*, Pearson, Spearman correlation coefficient.

**Figure S3.**
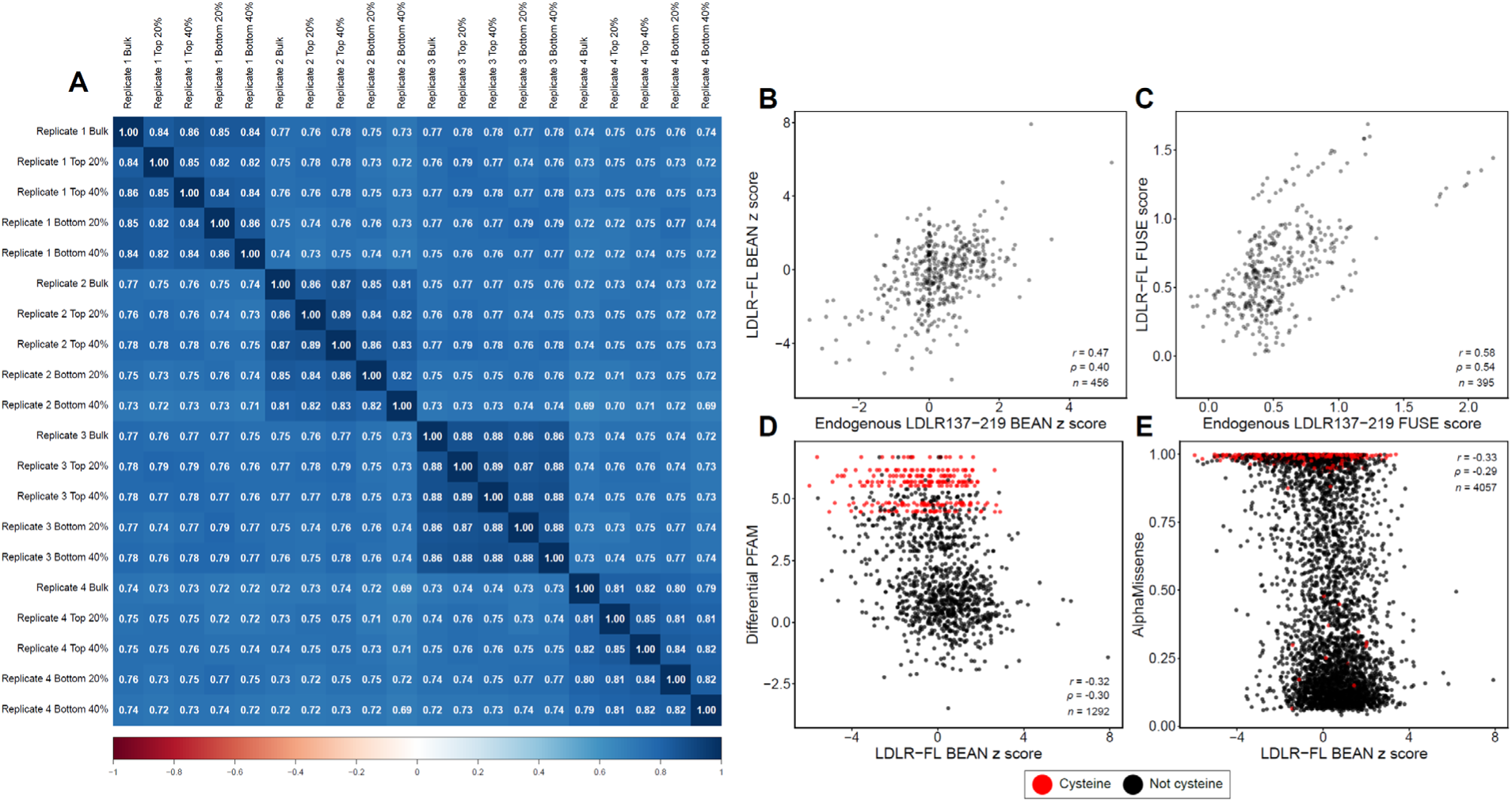
Assessing reproducibility and concordance of LDLR-FL data. **A,** Correlation matrix of technical replicates for sorted and bulk next-generation sequencing samples. **B-C,** Scatterplot of the endogenous LDLR137-219 versus LDLR-FL BEAN z-scores (**B**) and FUSE scores (**C**). **D-E**, Scatterplots of LDLR-FL BEAN z-scores versus Differential PFAM **(D**) and AlphaMissense (**E**) scores. *r*, *ρ*, Pearson, Spearman correlation coefficient.

**Figure S4.**
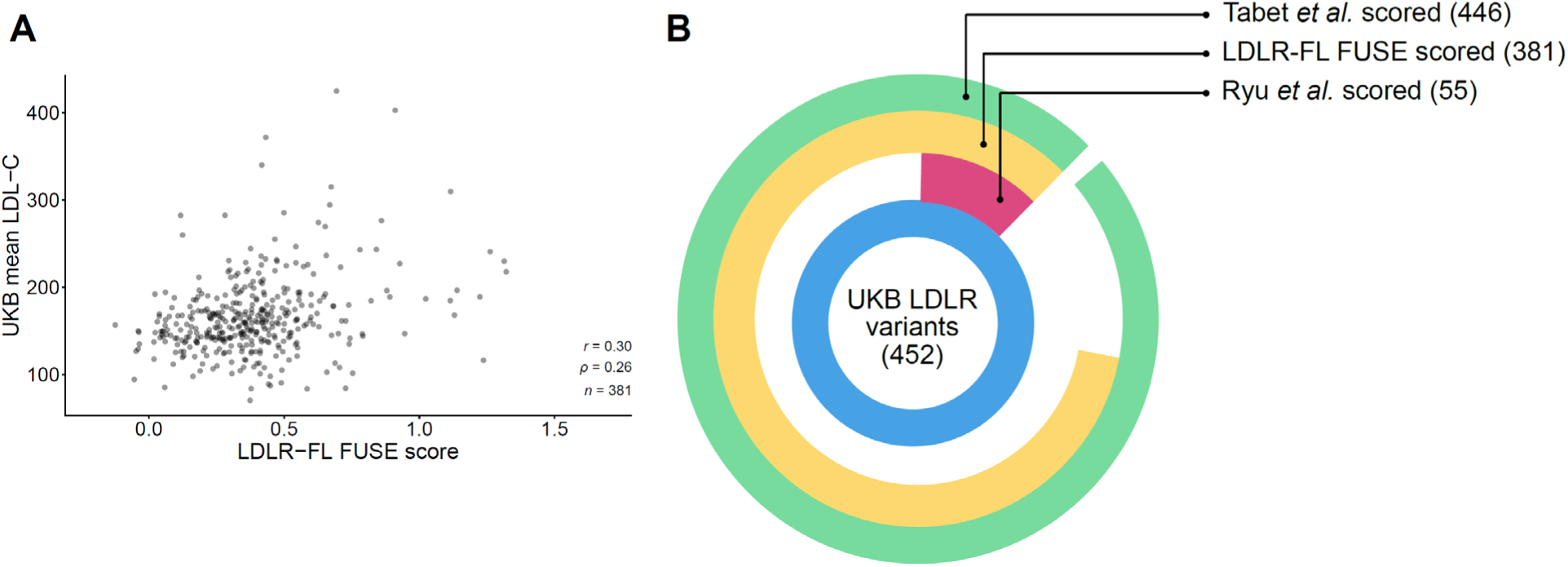
Comparison of LDLR-FL FUSE data to UK BioBank LDL-C levels. **A,** Scatterplot of LDLR-FL FUSE scores versus UK BioBank mean LDL-C levels. **B,** Pie chart of UK BioBank variant coverage by Ryu *et al.* 2024, LDLR-FL, and Tabet *et al.* 2025. *r*, *ρ*, Pearson, Spearman correlation coefficient.

**Figure S5.**
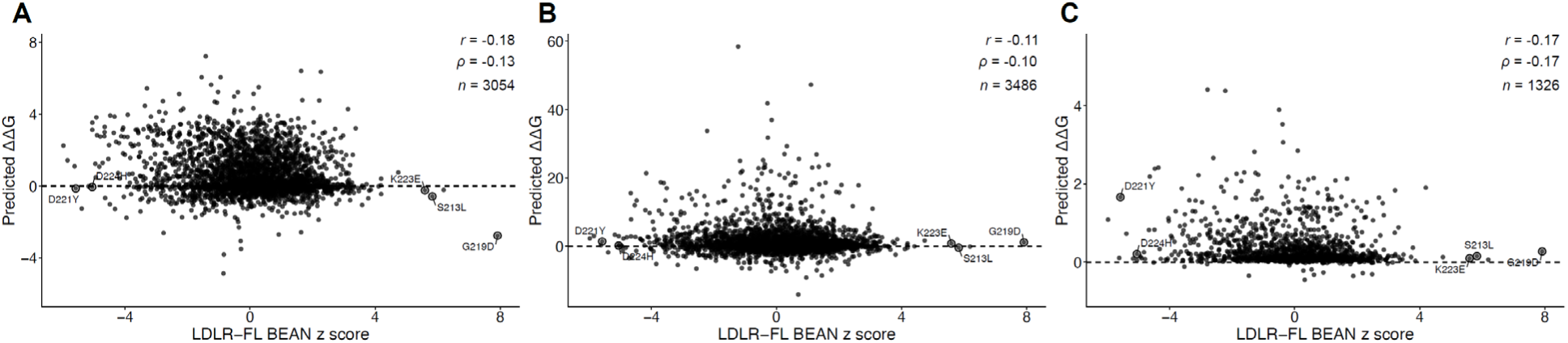
*In silico* stability and affinity analysis of LDLR variants. **A-C,** Scatterplots of LDLR-FL BEAN *z*-scores versus predicted ΔΔ*G* values from DDMut (**A**), FoldX (**B**), and DDMut-PPI (**C**). DDMut and FoldX predictions were run on the extracellular domain of LDLR (1n7d) and DDMut-PPI was computed on the LDLR class A repeats bound to the APOB beta-belt (9bde). Top variants from our LDLR-FL screen are circled in black. *r*, *ρ*, Pearson, Spearman correlation coefficient.

**Figure S6.**
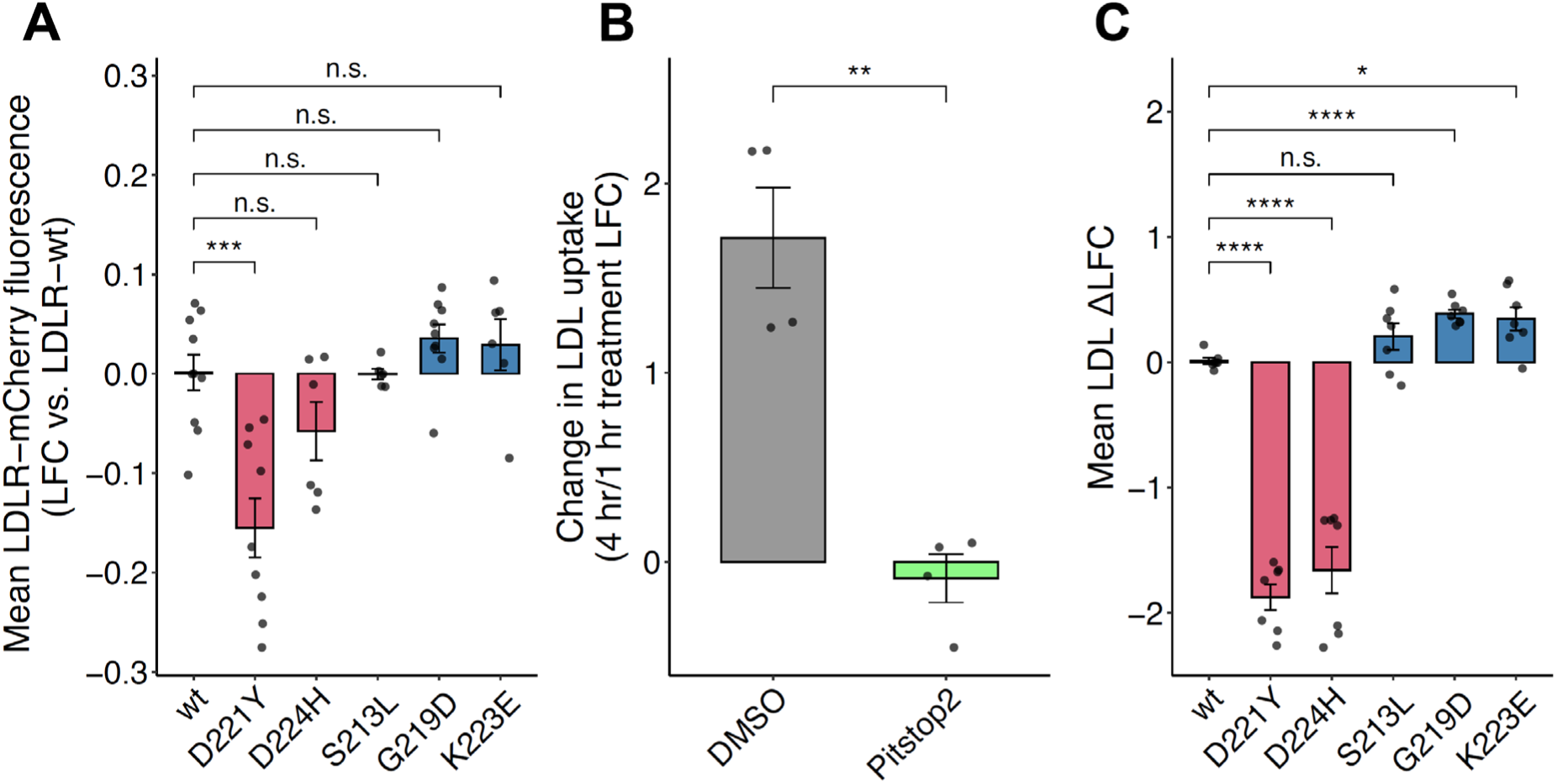
LDLR-mCherry expression and LDL binding versus uptake tests. **A,** Barplot of mean LDLR-mCherry fluorescence for single GOF and LOF variants. Wild type versus D221Y (*P* = 0.00056), wild type versus D224H (*P* = 0.12260), wild type versus S213L (*P* = 0.15278), wild type versus G219D (*P* = 0.39432), wild type versus K223E (*P* = 0.93655). **B,** Barplot of LDL uptake change in Pitstop2-treated versus DMSO-treated hepatocytes (*P* = 0.0028). **C,** Barplot of LDL uptake change in Pitstop2 treated hepatocytes for single GOF and LOF variants. Wild type versus D221Y (P = 6.8 x 10^-7), wild type versus D224H (P = 9.2 x 10^-5), wild type versus S213L (P = 0.118), wild type versus G219D (P = 2.2 x 10^-6), and wild type versus K223E (P = 0.011). \**P* ≤ 0.05, \*\**P* ≤ 0.01, \*\*\**P* ≤ 0.001, \*\*\*\**P* ≤ 0.0001; n.s., not significant (*P* > 0.05).

**Figure S7.**
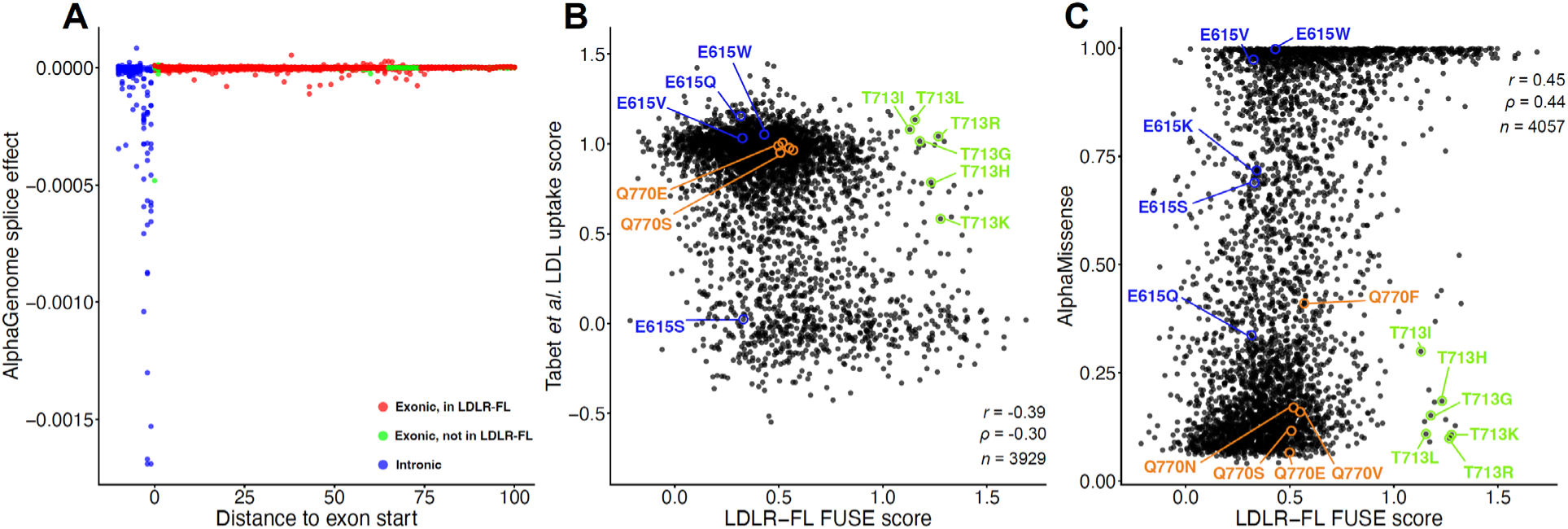
Variant effect characterization on LDLR splicing. **A,** AlphaGenome predicted splice effects of splice acceptor variants. **B-C,** Scatterplots of LDLR-FL FUSE scores versus Tabet *et al.* (**B**) and AlphaMissense (**C**) scores with variants at the three top splice donor sites circled in blue, orange, and green.

## Notes

### Competing Interest Statement

Luca Pinello has financial interests in Edilytics, Inc., and SeQure Dx, Inc. Gerald Schwank is a scientific advisor of Prime Medicine and co-founder of Nerai Bio. Authors' interests were reviewed and are managed by MassGeneral Brigham HealthCare in accordance with their conflict of interest policies.

